# Restoration of (nor)adrenergic-like Ca²⁺ signalling counteracts age-related motor decline

**DOI:** 10.64898/2026.09.28.754926

**Authors:** Anemari Horvat, Urška Černe, Anne-Kristin Dahse, Lara-Sophie Brodmerkel, Aljaž Bratanič, Madlen Matz-Soja, Stefanie Schirmeier, Robert Zorec, Nicole Scholz, Nina Vardjan

## Abstract

Aging disrupts neural function, but whether impaired noradrenergic control of brain metabolism contributes to functional decline remains unclear. Using genetically encoded fluorescent sensors we measured metabolic and signaling responses in neurons and glia of living *Drosophila* brains following stimulation with octopamine, the invertebrate analogue of noradrenaline. Aging was accompanied by neurodegeneration, reduced locomotion and enhanced oxidative whole-brain metabolism. Octopamine increased D-glucose uptake selectively in astrocytes and elevated L-lactate in both astrocytes and neurons, consistent with astrocytic glycolysis coupled to neuronal L-lactate uptake; monocarboxylate transporter inhibition caused selective astrocytic L-lactate accumulation. In aged animals octopamine-evoked metabolic responses in astrocytes and neural Ca²⁺ transients were markedly reduced. This impairment was associated with lower expression of adrenoceptor-like tyramine 1 receptor (Tyr_1_R). Selective Tyr_1_R overexpression in (nor)adrenergic-like Tdc2 neurons prolonged lifespan, restored Ca²⁺ signalling and improved locomotor performance. These findings identify Tyr_1_R-dependent Ca²⁺ signalling as an age-sensitive regulator of brain metabolism and motor function.

## Introduction

The nervous system is particularly vulnerable to aging, as age-associated neural changes increase susceptibility to neurodegenerative disorders, particularly Alzheimer’s and Parkinson’s disease ^1,2^. Research on brain aging has primarily focused on cognitive functions. However, aging also affects non-cognitive abilities, including visuomotor ^3^ and motor performance ^4,5^. Motor deficits strongly correlate with cognitive decline ^5^ and may even precede it ^4^.

A link between age-related decline in brain function and brain metabolism has been well established. One manifestation is reduced D-glucose utilization, i.e. hypometabolism, a hallmark of both physiological and pathological brain aging and a major risk factor for neurological decline. This metabolic imbalance drives neuronal, and likely also glial, dysfunction through aberrant mitochondrial activity and associated disruption in lipid metabolism, thereby further promoting aging or disease progression ^2,6,7^. Notably, metabolic alterations often precede the onset of clinical symptoms ^8,9^, highlighting brain metabolism as a promising therapeutic target. During periods of intense brain activity, the energy substrates D-glucose and L-lactate are supplied to neurons primarily by glia, while lipids (free fatty acids) are transported from neurons to glia ^10–12^. Thus, neuroglial metabolic coupling is key for normal brain function, and its disruption reflects in cognitive and behavioural deficits ^13^.

In mammals, brain metabolism is intricately modulated through the noradrenergic system ^14,15^. Noradrenergic projections that originate in a small nucleus in the brainstem (*locus coeruleus*), innervate virtually all brain regions. Noradrenaline binds to G protein-coupled adrenoceptors and activates intracellular Ca^2+^ and cAMP pathways to facilitate D-glucose uptake, glycogenolysis and aerobic glycolysis, which ultimately drive L-lactate production in adrenoceptor-rich astrocytes ^16^. According to the astrocyte-neuron L-lactate shuttle (ANLS) hypothesis, astrocyte-derived L-lactate serves as an energy substrate for neurons ^11^ and supports cognitive processes including learning and memory formation ^17–19^. Beyond its metabolic role, the noradrenergic system also shapes motoric ability through crosstalk with cognitive processes like attention, memory, and decision-making ^20^. Age-related dysfunction of noradrenergic signalling has been linked to impairment in both cognitive and motor function, highlighting this system as a suitable entry point for therapeutic intervention ^20–22^. However, it remains unclear whether noradrenergic malfunction causally contributes to the metabolic aberrations underlying age-related cognitive and motor decline, and whether these defects arise in a cell type-specific manner. Defining cell-specific targets in the brain could enable precise modulation of neuronal and glial function within the glia–neuron unit.

Here, we used *Drosophila melanogaster* as a model of aging to elucidate how age affects noradrenergic regulation of brain cell function, motoric performance and longevity. To this end, we analysed young (2–5 days) and aged flies (30–35 days) by virtue of locomotor performance, lifespan, neurodegeneration, and the responsiveness of neurons and glial cells in intact, living brains to octopamine (Oct), the analogue of mammalian noradrenaline, released from (nor)adrenergic-like tyrosine decarboxylase 2 (Tdc2) neurons. Genetically encoded fluorescent sensors were used to monitor Oct-induced changes in cytosolic D-glucose, L-lactate, cAMP, and Ca²⁺ selectively in central neurons and glia. Aged animals showed neurodegenerative signs, including pathological vacuoles, elevated oxidative brain metabolism and impaired motoric function. At the cellular level old age had several effects: First, the capacity for D-glucose uptake was diminished in neurons. Oct-induced increase in D-glucose occurred only in astrocytes and was attenuated with age. Second, while L-lactate was taken up by both neurons and astrocytes, this was not affected by age. Oct-facilitated glycolytic L-lactate production in astrocytes declined with age, whereas concomitant neuronal L-lactate uptake remained preserved. Third, independent of age, only neurons responded to Oct with cAMP changes.

Finally, neuronal and glial Ca²⁺ responses to Oct were almost completely abolished in aged brains. Overexpression of adrenergic-like tyramine 1 receptor (Tyr_1_R (Oct-TyrR)) in octopaminergic Tdc2-positive neurons reversed this phenotype to levels compareable to young animals. Consistent with these findings, at systemic levels overexpression of Tyr_1_R reversed age-related locomotor decline and even led to more than 20% increase in lifespan. Taken together, our findings suggest a model in which old age reduces Tyr_1_R expression and thereby abolishes Oct-dependent Ca^2+^ signalling in neurons and glia, with cell type-specific and systemic functional consequences. This work positions adrenoceptor-mediated Ca²⁺ signalling as a promising therapeutic target to interfere with neurological decline in aging and neurodegenerative disorders.

## Results

Aging will over time result in neurodegeneration and a decline in cognitive and motoric abilities ^2^. The aging brain undergoes changes in glucose metabolism (hypometabolism). As a key regulator of brain metabolism, cognition, and motor function ^23^, the noradrenergic system is thought to pose a contributing factor ^20,21^. However, if and how faulty noradrenergic regulation impacts age-dependent decline in cognition and motor performance remains unknown. To address this, we leveraged the short generation time, and experimental and genetic tractability of *Drosophila* to examine whether aging disrupts noradrenergic neural regulation ^1,24^. Here, we profiled the impact of old age on the metabolic and signalling capacity of neurons and different glial cells and identified adrenergic-like Tyr_1_R as a potential therapeutic entry point to reverse age-dependent decline on cellular and systemic levels.

### Aged flies show signs of neurodegeneration, neural metabolic changes and impaired locomotor performance

One of the common signs of neurodegeneration in the aging brain is an increase in the presence of pathological vacuoles in the brain parenchyma and brain atrophy ^25^. This structural deterioration is associated with altered brain energy metabolism, as functional neuroimaging studies with 2-deoxy-2-[^18^F]fluoro-D-glucose ([^18^F]FDG), show reduced brain D-glucose utilization (hypometabolism) ^8,9^, which correlates with impairments in cognitive function ^9^ and motor performance ^26^ in both humans and rodent models of aging and neurodegeneration. To establish aged *Drosophila* as a model to study aging, we first assessed whether brains of aged wild-type *Drosophila* (*w1118*) show this neurodegenerative trait. Using a protocol that preserves whole-brain architecture ^27^, brains from young (up to 5 days old) and aged (30–35 days old) flies were stained with the F-actin probe Alexa Fluor™ 647-conjugated phalloidin. Brain vacuolization, an indicator of neurodegeneration, quantified as the increase in number and area of pathological vacuoles ^27^, was more pronounced in aged than in young brains. In addition, the total vacuole area per brain was more than 2-fold higher in the brains of aged flies (Fig. 1A-B, Suppl. Table S2). These results are consistent with age-related neurodegeneration.

**Fig. 1:**
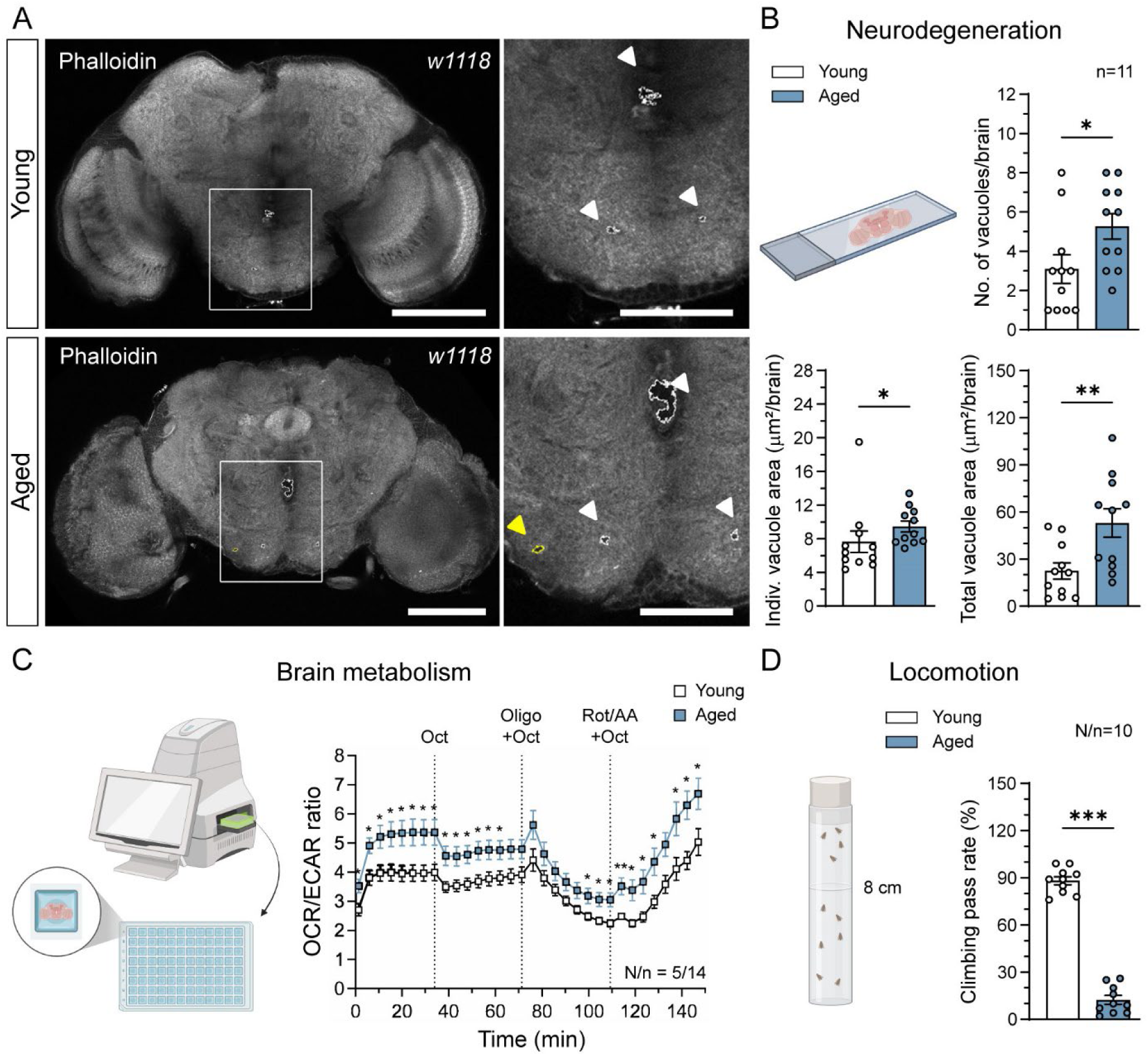
Aging in *Drosophila* increases the number and area of brain vacuoles, alters brain metabolism and reduces climbing capability. **A**) Representative single Z-stack images of fixed whole-brain mounts from young (upper panels) and aged (lower panels) wild-type *Drosophila* (*w1118*), labelled with Alexa Fluor^TM^ 647 Phalloidin (1:20). Images on the right represent white-framed areas on the left at higher magnification, showing physiologically present holes (white outlines/arrowheads) and vacuoles typical of neurodegeneration (yellow outline/arrowhead). Scale bars: 100 µm (left images), 50 µm (right images). **B**) Mean number of vacuoles (upper panel), individual vacuole area (in µm^2^; lower panel, left) and total vacuole area (in µm^2^; lower panel, right) per brain in young (white; n=11 brains) and aged (blue; n=11 brains) wild-type *Drosophila*. Data are presented as means ± SEM. Circles represent individual data points (mean values per brain). \**P* < 0.05, \*\**P* < 0.01 (Mann-Whitney *U*-test (No. of vacuoles and Indiv. vacuole area) or Unpaired t-test (Total vacuole area). **C)** Mean traces of OCR/ECAR ratio values over time under basal and stimulated conditions in brains from young (white) and aged (blue) wild-type (*w1118*) *Drosophila*. Brains were stimulated with 1 mM Oct, 10 µM oligomycin and 1 mM Oct (Oligo+Oct) and 5 µM rotenone/antimycin A and 1 mM Oct (Rot/AA+Oct) at 8^th^, 16^th^ and 24^th^ cycle, respectively (dotted lines). N=number of different runs (experiments), n=number of brains analyzed per age group. Each data point represents one measurement cycle. Data are presented as means ± SEM. \**P* < 0.05, \*\**P* < 0.01 (comparison of individual data points between young and aged brains; Unpaired t-test or Mann-Whitney *U*-test). **D**) Mean climbing pass rate (in %) of young (white) and aged (blue) wild-type *Drosophila*. N=number of independent vials (experiments), n=number of consecutive recordings per vial. Data are presented as means ± SEM. Circles represent individual data points (mean values per vial (independent experiment)). \*\*\**P* < 0.001 (Unpaired t-test). Graphical parts of figure were created in https://BioRender.com.

Aging and neurodegeneration are associated with metabolic changes in the brain ^9^. To explore this connection in our aging model, we measured the oxygen consumption rate (OCR) and extracellular acidification rate (ECAR) using a seahorse analyzer to gain insight into mitochondrial respiration and aerobic glycolysis rate, respectively, in young and aged living (*ex vivo*) wild-type brains (Fig. 1C). Under basal conditions, aged brains showed higher OCR/ECAR ratio than young brains, indicating a reduction in aerobic glycolysis with old age. When stimulated with the neuromodulator Oct, both age groups showed decreased OCR/ECAR (Fig. 1C), reflecting Oct-mediated facilitation of aerobic glycolysis and L-lactate release in young and aged brains.

Given that age-related neurodegeneration can negatively affect motoric ability ^28^, we assessed the motoric ‘fitness’ of young and aged wild-type flies in a negative geotaxis-based climbing assay ^29^. This analysis provides a combined measure of neuronal and muscular function ^28,29^. The climbing pass rate of old animals was more than 7-fold lower compared to young animals, with only ∼10% of aged animals successfully surpassing the 8-cm mark, compared to nearly ∼90% of young animals (Fig. 1D, Suppl. Table S2). These findings reveal a strong age-dependent decline in climbing performance.

In summary, our data show that aged animals develop neurodegenerative brain lesions, display aberrant brain metabolism, and exhibit a reduction in locomotor performance, establishing *Drosophila* as a compelling model for studying aging-related phenotypes.

### Aging selectively reduces neuronal D-glucose uptake, but does not affect L-lactate uptake

Given the age-dependent metabolic changes in whole brains (Fig. 1), we asked whether aging affects uptake of the essential metabolites D-glucose and L-lactate in specific cell types. We selectively expressed the FRET-based D-glucose sensor FLII^12^Pglu-700µδ6 in astrocytes (*alrm>FLII^12^Pglu-700µδ6*) or neurons (*elav>FLII^12^Pglu-700µδ6*) and measured cytosolic D-glucose levels in living brains (optic lobe) upon exposure to elevated D-glucose (Fig. 2A-D). Bath application of D-glucose (20 mM) elicited an immediate and sustained cytosolic free D-glucose rise, i.e., uptake, in astrocytes and neurons (Fig. 2A-B). Neurons of young brains showed a 1.4-fold (42.9%) greater (10.92 ± 1.08 vs. 7.64 ± 0.72; *P* < 0.05, Fig. 2C-D; Suppl. Table 4) and a 1.6-fold (38.9%) faster (Fig. 2I) relative D-glucose uptake than astrocytes. Importantly, aging had no effect on astroglial D-glucose uptake but decreased neuronal uptake by 42% (Fig. 2C-D, Suppl. Table S4).

**Fig. 2:**
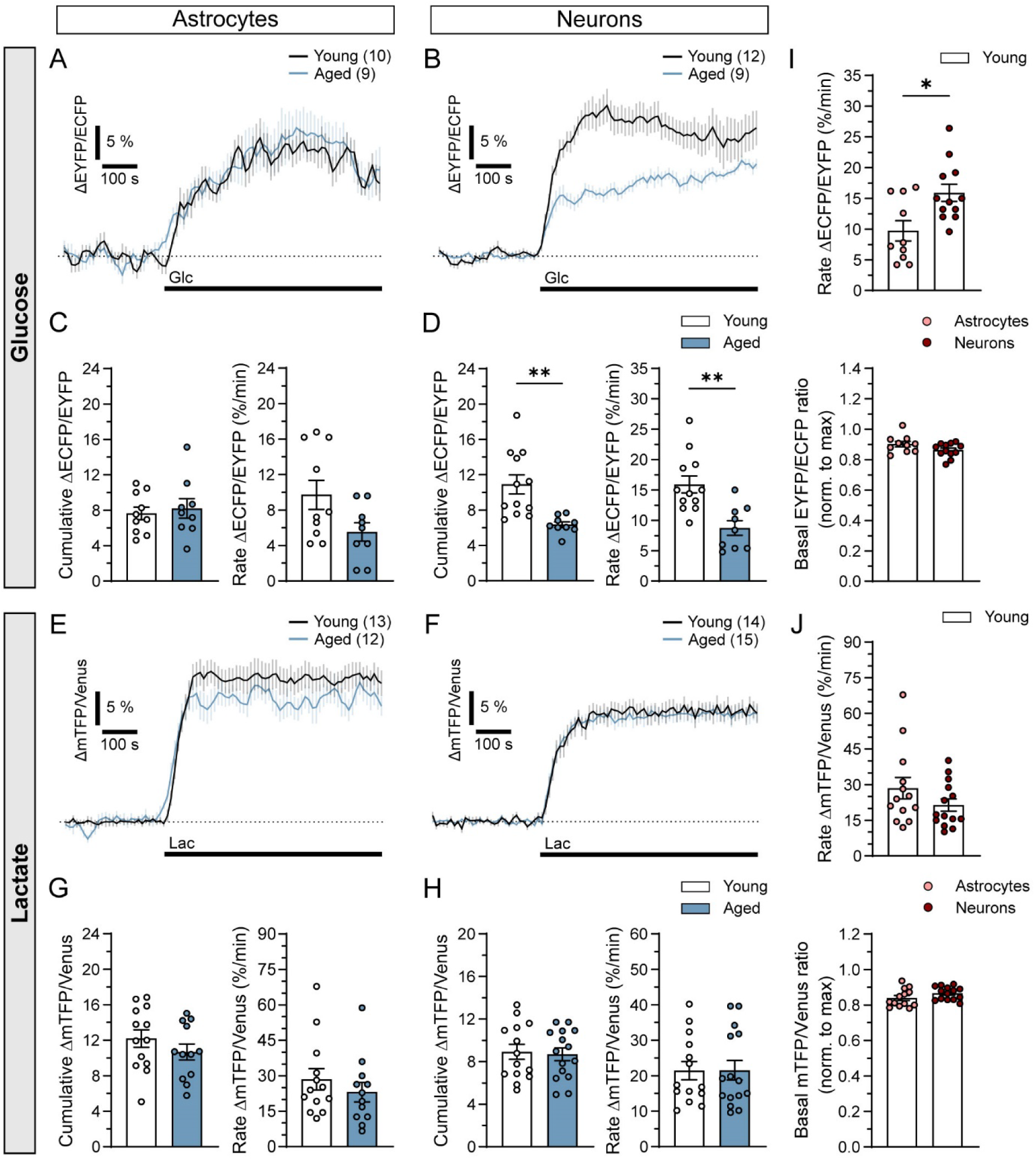
Exposure of *Drosophila* brains to elevated D-glucose or L-lactate triggers their accumulation in the cytosol of optic lobe astrocytes and neurons, with D-glucose accumulation in neurons being reduced in brains of aged *Drosophila*. **A-B)** Mean time-dependent changes in the FLII^12^Pglu-700µδ6 fluorescence ratio signal (ΔEYFP/ECFP) before and after the addition of 20 mM D-glucose (Glc; black lines) in A) astrocytes (*alrm-Gal4>UAS-FLII^12^Pglu-700µδ6*) and B) neurons (*elav-Gal4>UAS-FLII^12^Pglu-700µδ6*) in brains from young (black) and aged (blue) *Drosophila*. Data are expressed in percentages as a change in fluorescence ratio signal (ΔEYFP/ECFP) relative to the baseline values, reflecting changes in cytosolic D-glucose levels. Numbers in parentheses depict the number of brains analyzed. **C-D)** Mean cumulative changes (Cumulative ΔEYFP/ECFP; left) and rate of changes in the FLII^12^Pglu-700µδ6 ratio signal (Rate ΔEYFP/ECFP (%/min); right) after the addition of 20 mM Glc in C) astrocytes and D) neurons in brains from young (white) and aged (blue) *Drosophila*. Data are presented as means ± SEM. Circles represent individual data points (mean values of all analyzed astrocytes or neurons within individual optic lobe). \*\**P* < 0.01 (Unpaired t-test (D, right) or Unpaired t-test with Welch’s correction (D, left)). **E-F)** Mean time-dependent changes in the Laconic fluorescence ratio signal (ΔmTFP/Venus) before and after the addition of 30 mM L-lactate (Lac; black lines) in E) astrocytes (*alrm-Gal4>UAS-Laconic*) and F) neurons (*elav-Gal4>UAS-Laconic*) in brains from young (black) and aged (blue) *Drosophila*. Data are expressed in percentages as a change in fluorescence ratio signal (ΔmTFP/Venus) relative to the baseline values, reflecting changes in cytosolic L-lactate levels. Numbers in parentheses depict the number of brains analyzed. **G-H)** Mean cumulative changes (Cumulative ΔmTFP/Venus; left) and rate of changes in the Laconic ratio signal (Rate ΔmTFP/Venus (%/min); right) after the addition of 30 mM Lac in G) astrocytes and H) neurons in brains from young (white) and aged (blue) *Drosophila*. Data are presented as means ± SEM. Circles represent individual data points (mean values of all analyzed astrocytes or neurons within individual optic lobe). **I-J)** Mean rate of changes in the I) FLII^12^Pglu-700µδ6 (Rate ΔEYFP/ECFP (%/min); upper panel) and J) Laconic ratio signal (Rate ΔmTFP/Venus (%/min); upper panel) and mean baseline values of I) FLII^12^Pglu-700µδ6 (EYFP/ECFP; lower panel) and J) Laconic (mTFP/Venus; lower panel) fluorescence ratio signal normalized to maximal ratio changes upon stimulation with 20 mM Glc and 30 mM Lac, respectively, reflecting basal cytosolic D-glucose and L-lactate levels, in astrocytes and neurons in brains from young *Drosophila*. Data are presented as means ± SEM. Circles represent individual data points (mean values of all analyzed astrocytes or neurons within individual optic lobe). \**P* < 0.05 (Mann-Whitney *U*-test).

Next, we examined the impact of age on extracellular L-lactate uptake. We expressed the L-lactate FRET sensor Laconic selectively in astrocytes (*alrm>Laconic*) or neurons (*elav>Laconic*) and measured cytosolic L-lactate levels in optic lobes of young and aged brains upon bath application of high extracellular L-lactate (30 mM; Fig. 2E-H). When exposed to L-lactate, both astrocytes and neurons showed an immediate and sustained, age-independent elevation in cytosolic L-lactate (Fig. 2E-F), indicating L-lactate uptake. Age had no effect on the L-lactate uptake in either astrocytes or neurons (Fig. 2G-H; Suppl. Table S5).

Similar to *Drosophila* astrocytes also in cortex glia, another *Drosophila* glial cell type with analogous function to mammalian astrocytes, we observed no age-related differences in D-glucose or L-lactate uptake (Suppl. Fig. S1, Suppl. Tables S4 and S5).

Basal D-glucose and L-lactate levels were similar between astrocytes and neurons in young brains (Fig. 2I-J, lower panels), whereas in aged brains neuronal D-glucose levels were significantly lower than in astrocytes (Suppl. Fig. S2; Suppl. Table S3), consistent with the observed age-related reduction in extracellular D-glucose uptake in neurons (Fig. 2B).

These results show that age does not affect L-lactate uptake, but it reduces basal neuronal D-glucose levels relative to astrocytes and impairs D-glucose uptake, suggesting age-related alterations in neuronal D-glucose transport mechanisms.

### Octopamine evokes an astrocyte-specific rise in cytosolic D-glucose that is lost with age

In *Drosophila*, Tdc2-positive neurons exert metabolic control through adrenergic-like Oct/Tyr receptors ^30,31^. To identify the cell type responsible for age-related metabolic alterations in *Drosophila* brains and locomotor deficit (Fig. 1), we examined the impact of aging on Oct-mediated modulation of neuronal and glial D-glucose metabolism. To this end, cytosolic D-glucose levels were quantified in optic lobe astrocytes (*alrm>FLII^12^Pglu-700µδ6*), cortex glia (*R55B12>FLII^12^Pglu-700µδ6*) or neurons (*elav>FLII^12^Pglu-700µδ6*) in young and aged living brains following Oct bath application (Fig. 3).

**Fig. 3:**
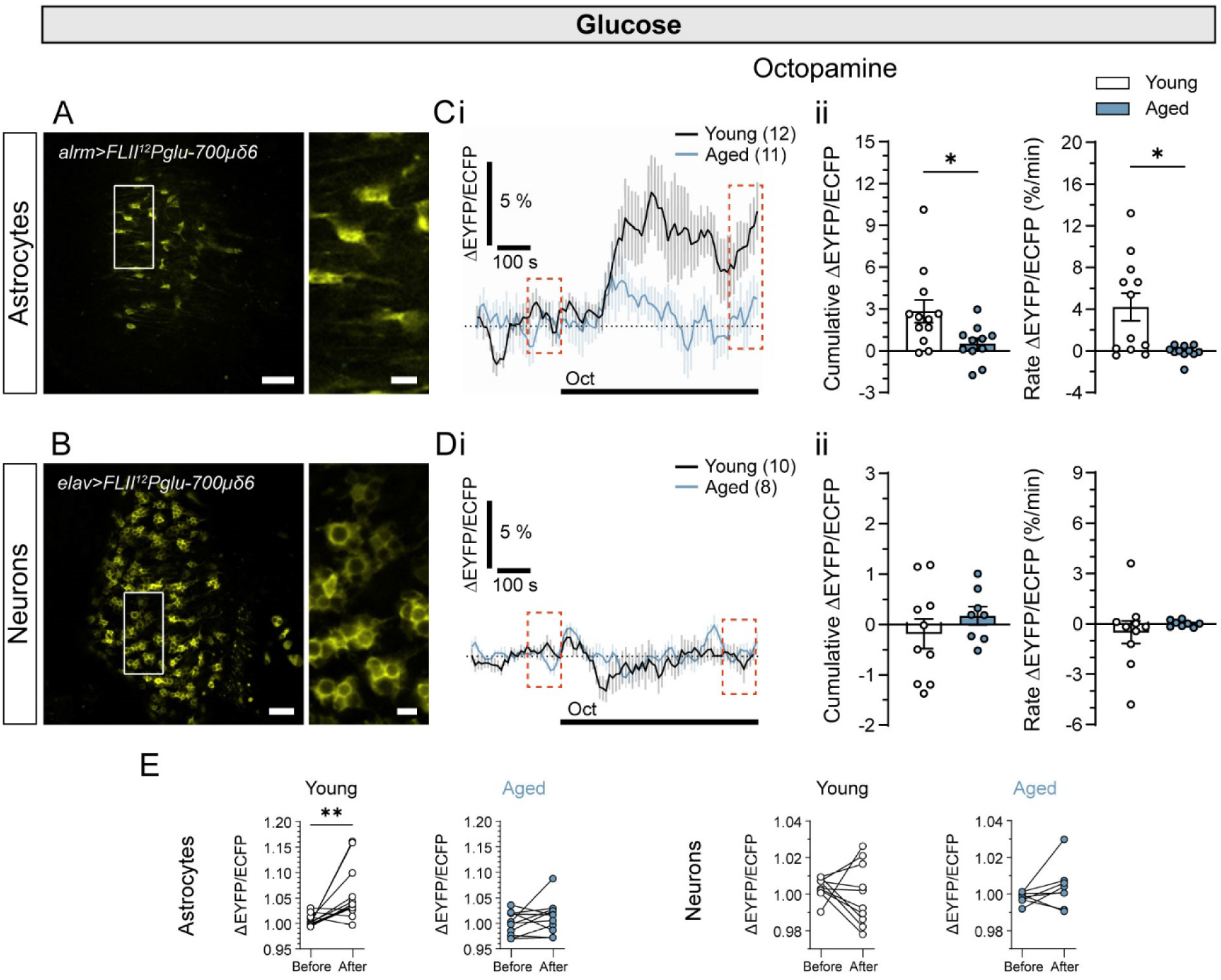
Octopamine induces sustained cytosolic D-glucose elevations in astrocytes, but not in neurons, in the optic lobes of young *Drosophila* brains only. **A-B)** Representative fluorescence images of brain optic lobes of young *Drosophila* expressing the D-glucose FRET sensor FLII^12^Pglu-700µδ6 in A) astrocytes (*alrm>FLII^12^Pglu-700µδ6*) and B) neurons (*elav>FLII^12^Pglu-700µδ6*). Images show merged EYFP and ECFP fluorescence emissions of the FRET sensor. Right panels show white framed regions on the left at higher magnification. Scale bars: 20 μm and 5 μm (insets). **Ci-Di)** Mean time-dependent changes in the FLII^12^Pglu-700µδ6 fluorescence ratio signal (ΔEYFP/ECFP) before and after the addition of 10 µM octopamine (Oct; black lines) in Ci) astrocytes and Di) neurons in brains from young (black) and aged (blue) *Drosophila*. Data are expressed as percentages of change in fluorescence ratio signal (ΔEYFP/ECFP) relative to baseline values, reflecting changes in cytosolic D-glucose levels. Numbers in parentheses depict the number of brains analyzed. **Cii-Dii)** Mean cumulative changes (Cumulative ΔEYFP/ECFP; left) and rate of changes in the FLII^12^Pglu-700µδ6 ratio signal (Rate ΔEYFP/ECFP (%/min); right) after the addition of 10 μM Oct in Cii) astrocytes and Dii) neurons in brains from young (white) and aged (blue) *Drosophila*. Data are presented as means ± SEM. Circles represent individual data points (mean values of all analyzed astrocytes or neurons within individual optic lobe). \**P* < 0.05 (Mann-Whitney *U*-test). **E)** Paired mean ratio signal values of 100 s before stimulation and mean ratio signal values of the last 100 s upon stimulation with 10 µM Oct in astrocytes and neurons in brains of young and aged *Drosophila* (indicated by red dotted squares in Ci and Di). \*\**P* < 0.001 (Wilcoxon test).

In young brains, Oct stimulation elicited a sustained increase in cytosolic D-glucose in astrocytes (Fig. 3C), likely due to OctR/TyrR-mediated ^32^ facilitation of extracellular D-glucose uptake ^16^. The increase occurred with a ∼100 s delay after Oct application, further suggesting a receptor-mediated response that may involve activation of intracellular signalling pathways before the metabolic response becomes apparent. The facilitation of D-glucose uptake was reduced by 82% in older brains (Fig. 3C and E; Suppl. Table S4). Oct-dependent D-glucose uptake was not detected in neurons or cortex glia. This remained true in aged brains as well (Fig. 3D and E and Suppl. Figs. S3 and S6, respectively; Suppl. Table S4).

Together, these data show that Oct induces astrocyte-specific facilitation of D-glucose uptake in the *Drosophila* brain, which is lost with age.

### Octopamine drives sustained cytosolic L-lactate rise in neurons and astrocytes, with weaker and age-declining responses in astrocytes

To examine the impact of aging on Oct-modulated aerobic glycolysis in the *Drosophila* brain cells, we monitored cytosolic L-lactate responses to Oct using the genetically encoded Laconic sensor. Laconic was expressed selectively in astrocytes (*alrm>Laconic*) or neurons (*elav>Laconic*), and Oct-induced changes in cytosolic L-lactate were recorded in the optic lobes of young and aged brains by real-time confocal microscopy (Fig. 4).

**Fig. 4:**
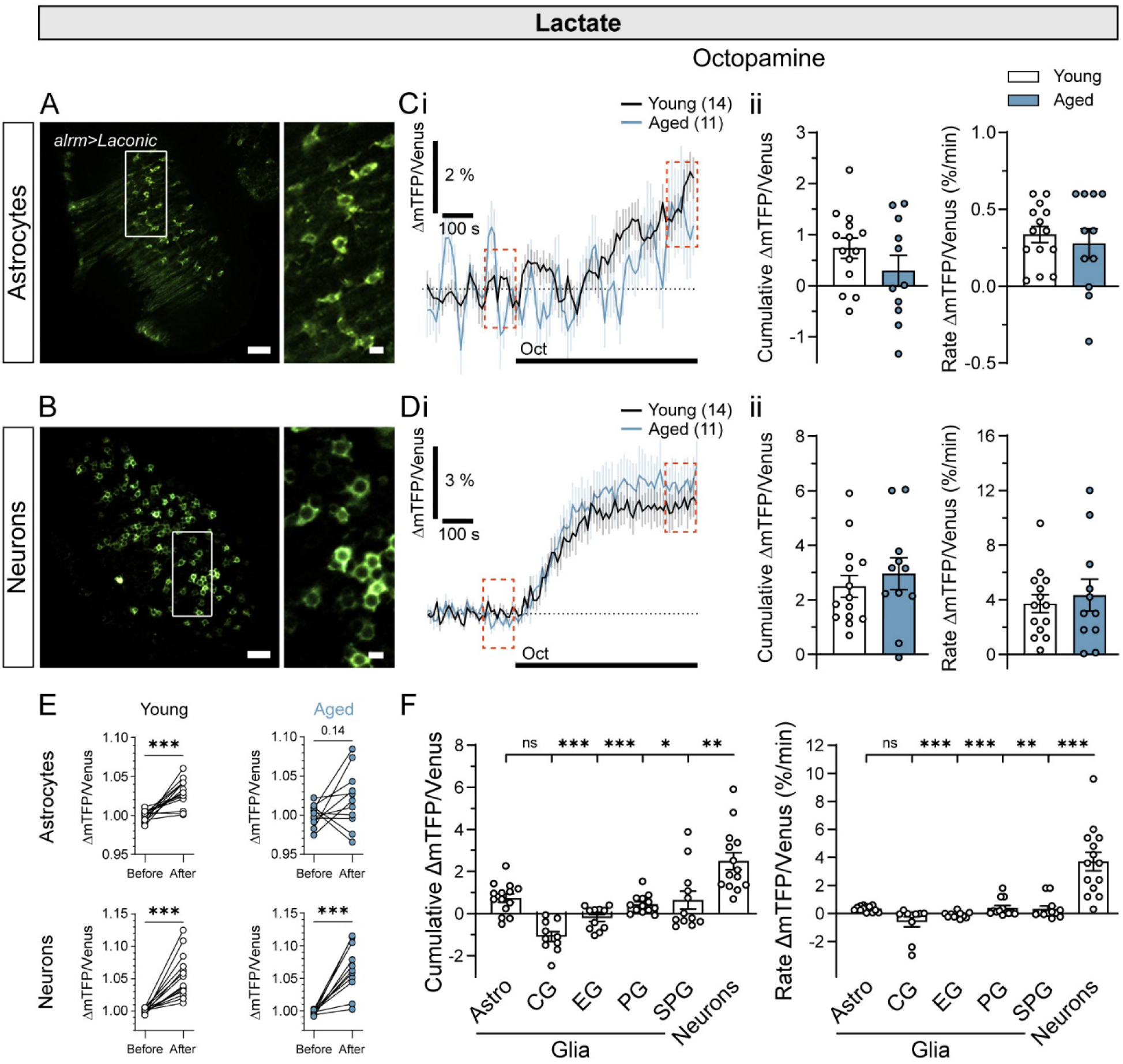
Octopamine induces sustained cytosolic L-lactate elevations in astrocytes and neurons in the *Drosophila* brain optic lobes, which are in astrocytes less pronounced and reduced with aging. **A-B)** Representative fluorescence images of brain optic lobes of young *Drosophila* expressing the L-lactate FRET sensor Laconic in A) astrocytes (*alrm>Laconic*) and B) neurons (*elav>Laconic*). Images show merged mTFP and Venus fluorescence emissions of the FRET sensor. Right panels show white-framed regions on the left at higher magnification. Scale bars: 20 µm and 5 µm (insets). **Ci-Di)** Mean time-dependent changes in the Laconic fluorescence ratio signal (ΔmTFP/Venus) before and after the addition of 10 µM octopamine (Oct; black lines) in Ci) astrocytes and Di) neurons in brains from young (black) and aged (blue) *Drosophila*. Data are expressed as percentages of change in fluorescence ratio signal (ΔmTFP/Venus) relative to baseline values, reflecting changes in cytosolic L-lactate levels. Numbers in parentheses depict the number of brains analyzed. **Cii-Dii)** Mean cumulative changes (Cumulative ΔmTFP/Venus; left) and rate of changes in the Laconic ratio signal (Rate ΔmTFP/Venus (%/min); right) after the addition of 10 µM Oct in Cii) astrocytes and Dii) neurons in brains from young (white) and aged (blue) *Drosophila*. **E)** Paired mean ratio signal values of 100 s before stimulation and mean ratio signal values of the last 100 s upon stimulation with 10 µM Oct in astrocytes and neurons in brains of young and aged *Drosophila* (indicated by red dotted squares in Ci). \*\*\**P* < 0.001 (Paired t-test). **F**) Mean cumulative changes (Cumulative ΔmTFP/Venus; left) and rate of changes in the Laconic ratio signal (Rate ΔmTFP/Venus (%/min); right) after the addition of 10 µM Oct in astrocytes (Astro), cortex glia (CG; *R55B12>Laconic*), neuropil ensheathing glia (EG; *R83E12>Laconic*), perineurial glia (PG; *R85G01>Laconic*), subperineurial glia (SPG; *SPG>Laconic*) and neurons in young *Drosophila* brains. Data are presented as means ± SEM. Circles represent individual data points (mean values of all analyzed cells or region(s) within individual optic lobe/brain). \**P* < 0.05, \*\**P* < 0.01, \*\*\**P* < 0.001 (One-way ANOVA on ranks, followed by Dunn’s test).

Oct triggered L-lactate increases in astrocytes and neurons (Fig. 4C, D and E). In young *Drosophila* brains, inhibition of monocarboxylate transporters (MCTs) with α-cyano-4-hydroxycinnamic acid (CHC) resulted in an approximately 4-fold higher rate of L-lactate accumulation in astrocytes than in neurons (Suppl. Fig. S4; Suppl. Table S5), indicating substantially greater intrinsic L-lactate production, and thus aerobic glycolytic activity, in astrocytes. Consistent with this, in mammalian systems, astrocytes have a greater capacity to produce L-lactate through aerobic glycolysis than neurons, ^11^. These findings suggest that the Oct-evoked increase in astrocytic L-lactate primarily reflects stimulation of aerobic glycolysis, whereas the increase observed in neurons is more likely driven by uptake of extracellular L-lactate.

In aged astrocytes, the mean cumulative change and rate of change in the Laconic signal upon Oct stimulation did not differ from those in young astrocytes (Fig. 4Cii; Suppl. Table S5). Yet, the mean change in Laconic signal relative to baseline (Fig. 4E) and to vehicle stimulation (Suppl. Fig. S6 and Suppl. Table S8) was no longer statistically significant in aged animals, which suggests a reduction in Oct-induced aerobic glycolysis in astrocytes with age. Contrarily, age did not affect Oct-induced L-lactate responses in neurons (Fig. 4D and E).

To examine if Oct affects cytosolic L-lactate in other glial cell types, we assessed Laconic responses towards Oct exposure selectively in cortex glia (*R55B12>Laconic*), neuropil ensheathing glia (*R83E12>Laconic*), perineurial glia (*R85G01>Laconic*) and subperineurial glia (*SPG>Laconic*). All cell types were unaffected, except for astrocytes. Compared to neurons, however, the mean cumulative and rate of change in the Laconic signals were significantly lower in cortex glia, neuropil ensheathing glia, perineurial glia and subperineurial glia (Fig. 4F). In the case of cortex glia, we also tested whether aging affects Oct-related cytosolic L-lactate levels, but no changes were observed in young vs. aged flies (Suppl. Figs. S5 and S6; Suppl. Tables S5 and S8).

In sum, we demonstrate that Oct evokes cytosolic L-lactate increases most prominently in neurons, independent of age, likely reflecting enhanced L-lactate uptake. In astrocytes, the Oct-induced L-lactate increase is overall lower, reflecting facilitation of aerobic glycolysis, which diminishes with age. In other glial cell types, Oct has no impact on cytosolic L-lactate.

### Octopamine triggers sustained cAMP increase in neurons irrespective of age

Oct modulates intracellular G_q_/Ca²⁺, G_s_/cAMP or G_i_/cAMP pathways through adrenergic-like Oct/Tyr receptors ^33,34^. Thus, we hypothesized that the aberrant Oct-mediated D-glucose and L-lactate metabolism in aged brains (Figs. 3 and 4) arises from age-related alterations in OctR/TyrR expression and thereby corrupts intracellular Ca^2^⁺ and/or cAMP signalling. To test this, we investigated the effect of Oct on cAMP levels in optic lobe astrocytes or neurons using the cAMP sensor Epac1-camps (*alrm>Epac1-camps* or *elav>Epac1-camps,* respectively) and assessed whether age played a role (Fig. 5). Application of Oct resulted in exponential rise in cytosolic cAMP in neurons irrespective of age. In contrast, astrocytes and cortex glia did not respond to Oct with cytosolic cAMP increases in either age group (Fig. 5C-D; Suppl. Figs. S7 and S9; Suppl. Tables S6 and S8). Our data show that Oct elicits a neuron-specific and age-independent sustained cytosolic rise in cAMP.

**Fig. 5:**
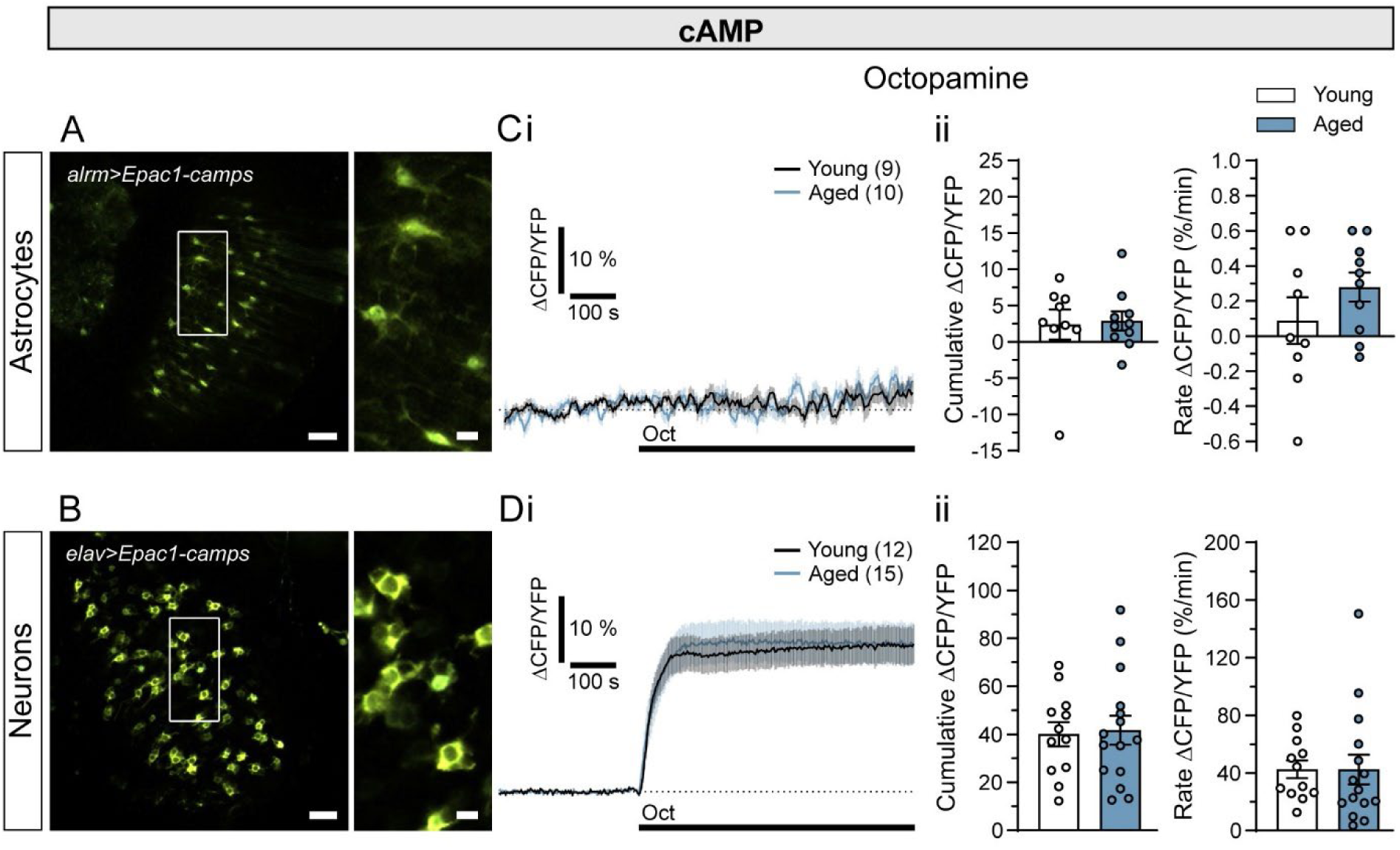
Octopamine induces sustained cytosolic cAMP elevations in neurons, but not in astrocytes, in the brain optic lobes of young and aged *Drosophila*. **A-B)** Representative fluorescence images of brain optic lobes of young *Drosophila* expressing the cAMP FRET sensor Epac1-camps in A) astrocytes (*alrm>Epac1-camps*) and B) neurons (*elav>Epac1-camps*). Images show merged CFP and YFP fluorescence emissions of the FRET sensor. Right panels show white-framed regions on the left at higher magnification. Scale bars: 20 µm and 5 µm (insets). **Ci-Di)** Mean time-dependent changes in the Epac1-camps fluorescence ratio signal (ΔCFP/YFP) before and after the addition of 10 µM octopamine (Oct; black lines) in Ci) astrocytes and Di) neurons in brains from young (black) and aged (green) *Drosophila*. Data are expressed as percentages of change in the ΔCFP/YFP signal relative to baseline values, reflecting changes in cytosolic concentration of cAMP. Numbers in parentheses depict the number of brains analyzed. **Cii-Dii)** Mean cumulative changes (Cumulative ΔCFP/YFP; left) and rate of changes in the Epac1-camps ratio signal (Rate ΔCFP/YFP (%/min); right) after the addition of 10 μM Oct in Cii) astrocytes and Dii) neurons in brains from young (white) and aged (green) *Drosophila*. Data are presented as means ± SEM. Circles represent individual data points (mean values of all analyzed astrocytes or neurons within individual optic lobe).

### Octopamine triggers cytosolic Ca^2+^ transients in astrocytes and neurons, which fade with age

Next, we investigated the effect of age on Oct-induced Ca^2+^ signaling in different brain cells. We used *Drosophila* lines expressing Ca^2+^ sensor jGCaMP7b selectively in astrocytes (*alrm>jGCaMP7b;mCherry*) and neurons (*elav>jGCaMP7b;mCherry*) and measured cytosolic Ca^2+^ signals of young and old optic lobe cells in response to bath application of Oct ^32^ (Fig. 6).

**Fig. 6:**
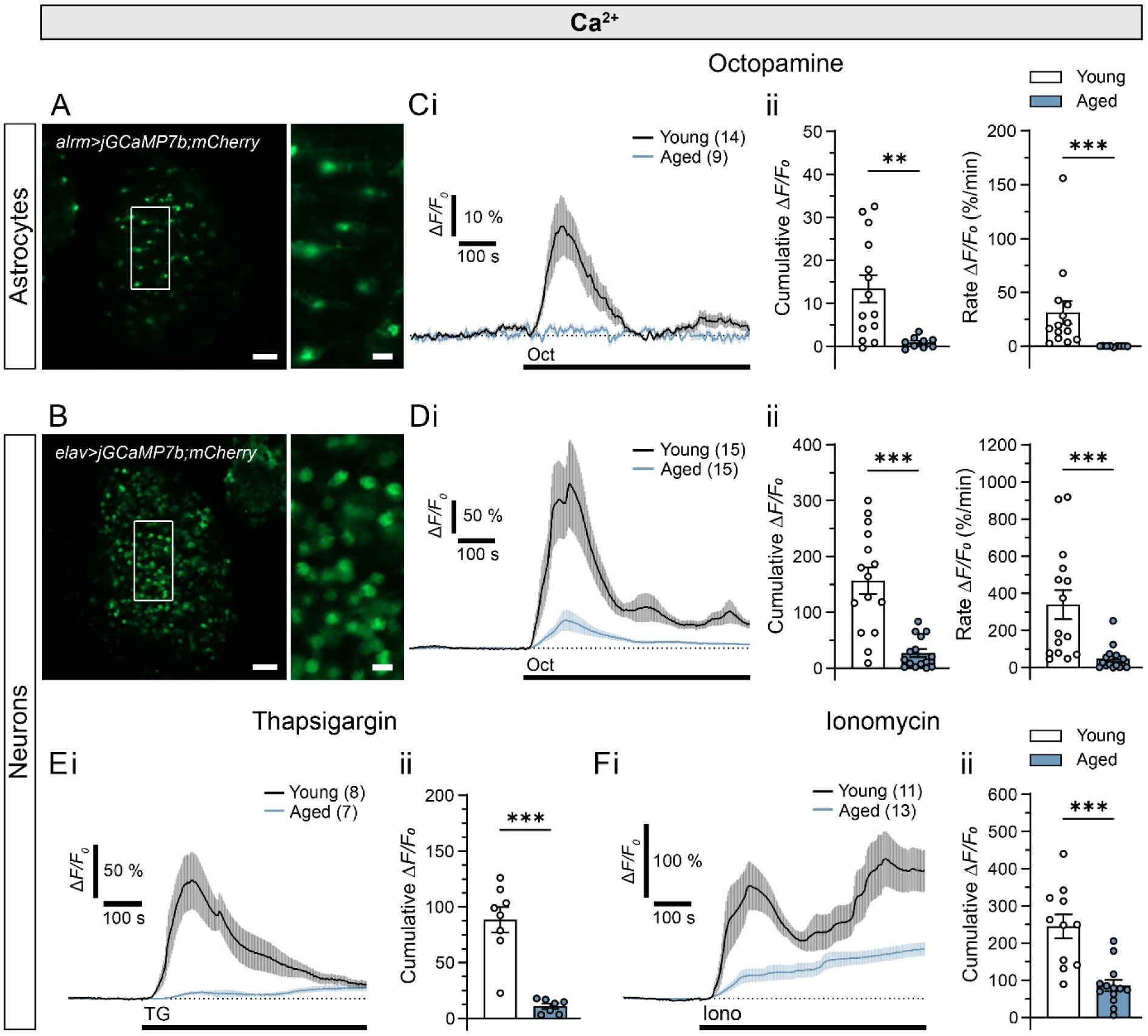
Octopamine-induced cytosolic Ca_2+_ transients are attenuated in astrocytes and neurons in the brain optic lobes of aged *Drosophila*. **A-B)** Representative fluorescence images of brain optic lobes of young *Drosophila* expressing the Ca^2+^ sensor jGCaMP7b in A) astrocytes (*alrm>jGCaMP7b;mCherry*) and B) neurons (*elav>jGCaMP7b;mCherry*). Right panels show white-framed regions on the left at higher magnification. Scale bars: 20 µm and 5 µm (insets). **Ci-Fi)** Mean time-dependent changes in jGCaMP7b fluorescence intensity signal (Δ*F/F_0_*) before and after the addition of Ci-Di) 10 µM octopamine (Oct; black lines) in Ci) astrocytes and Di) neurons and before and after the addition of Ei) 1 µM thapsigargin (TG) and Fi) 10 µM ionomycin (Iono; black lines) in neurons in brains from young (black) and aged (blue) *Drosophila*. Data are expressed in percentages as a change in the Δ*F/F_0_* signal relative to the baseline values, reflecting changes in cytosolic concentration of Ca^2+^. Numbers in parentheses depict the number of brains analyzed. **Cii-Fii)** Mean cumulative changes (Cumulative Δ*F/F_0_*; left) and rate of changes in the jGCaMP7b Δ*F/F_0_* signal (Rate Δ*F/F_0_* (%/min); right) after the addition of 10 µM Oct in Cii) astrocytes and Dii) neurons and after the addition of Eii) 1 µM TG and Fii) 10 µM Iono in neurons in brains from young (white) and aged (blue) *Drosophila*. Data are presented as means ± SEM. Circles represent individual data points (mean values of all analyzed astrocytes or neurons within individual optic lobe). \*\**P* < 0.01, \*\*\**P* < 0.001 (Mann-Whitney *U*-test (Cii (right) and Dii) or Unpaired t-test with Welch’s correction (Cii (left), Eii and Fii)).

In young brains, Oct induced a robust, transient increase in cytosolic Ca^2+^ in optic lobe astrocytes and neurons. This response is likely mediated through Oct activation of G_q_-coupled Oct/Tyr receptors ^32^. Strikingly, Oct-dependent Ca^2+^ transients were completely abolished in both neurons and astrocytes in aged brains (Fig. 6Ci-Di). In astrocytes, the mean cumulative change and rate of change in the jGCaMP7b signal after the addition of Oct were ∼14-fold and ∼200-fold lower in brains from aged compared to young *Drosophila*, respectively (Fig. 6Cii; Suppl. Table S7). In neurons, the mean cumulative change and the rate of change in the jGCaMP7b signal in response to Oct were ∼6-fold and ∼7-fold lower in brains from aged compared to young flies, respectively (Fig. 6Dii; Suppl. Table S7). Similar age-dependent attenuation of Oct-induced cytosolic Ca^2+^ transients was observed in cortex glia (*R55B12>jGCaMP7b;mCherry*; Suppl. Fig. S8; Suppl. Table S7).

These findings uncover an age-dependent impairment in Oct-mediated mobilisation of cytosolic Ca²⁺ in both glial cells and neurons of the *Drosophila* brain.

### Age reduces the capacity for neuronal Ca^2+^ mobilization

To investigate how age affects Ca^2+^ signaling, we first quantified the expression levels of the synaptic voltage-gated Ca^2+^ channel Cacophony ^35,36^. Visualisation of GFP tagged Cacophony expressed from the native promotor showed no overt difference in young vs. aged brains (^37^; Suppl. Fig. S10; Suppl. Table). Next, we exposed jGCaMP7b-expressing neurons of young and aged brains to TG (1 µM), a SERCA pump inhibitor that prevents Ca^2+^ reuptake into the ER ^38^, and to Iono (10 µM), a ionophore that promotes Ca^2+^ transport across cellular membranes ^39^, and monitored Ca^2+^ responses. Cytosolic Ca^2+^ increases following stimulation with either TG or Iono were significantly lower in aged neurons compared to young ones (Fig. 6E-F; Suppl. Table S7). This suggests that aging alters the storage capacity of intracellular Ca^2+^ and the efficacy of Ca^2+^ transport across neuronal membranes. Collectively, these changes may explain the reduction in the cell’s capacity for intracellular Ca^2+^ mobilisations.

### Aging leads to downregulation of adrenergic-like G_q_-coupled tyramine 1 receptor in *Drosophila* brain

Octopamine is sensed at the cell surface, which begs the question whether Oct receptors that utilize G_q_/Ca^2+^ pathways are involved in age-dependent changes. To test this, we examined the expression pattern of adrenergic-like G_q_-coupled Tyr_1_R in young and aged *Drosophila* brains (Fig. 7A). For this, we used a translational Tyr_1_R::T2A::GAL4 reporter to expresses Tyr_1_R alongside the yeast transcription factor GAL4 that drives expression of UAS-Transtimer ^40^, i.e., a fast-folding GFP (sfGFP) and a slow-maturing RFP selectively in Tyr_1_R-positive cells. GAL4 expression scales with Tyr_1_R quantity enabling insight into endogenous Tyr_1_R expression status at different time points. The mean fluorescence intensity ratio between sfGFP and RFP (sfGFP/RFP) served as a proxy for Tyr_1_Rs expression levels and uncovered a 30% reduction in Tyr_1_R expression in the central brain of aged animals (Fig. 7A; Suppl. Table S11). This suggests that Tyr_1_R expression declines with age and likely contributes to the dysregulation of Oct-dependent Ca^2+^ signalling in neural cells.

**Fig. 7:**
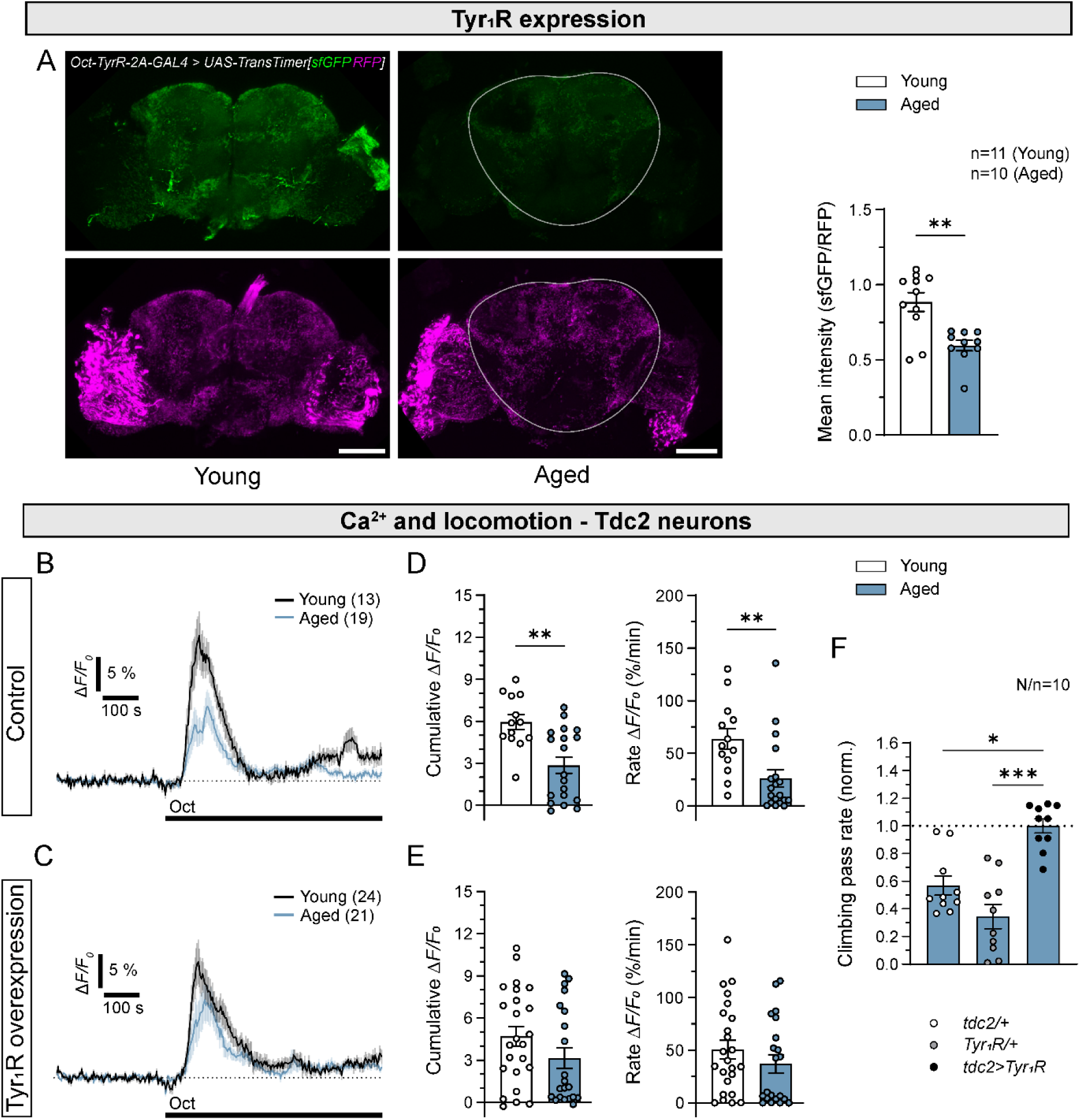
Overexpression of tyramine 1 receptors in Tdc2 neurons improves climbing capability of aged flies and recovers impaired Ca^2+^ response to octopamine in Tdc2 neurons in the brain optic lobes of aged *Drosophila*. **A)** Representative fluorescence images of young (left) and aged (right) *Drosophila* brains expressing a fast-folding GFP (sfGFP; green) and a slow-folding RFP (magenta) under the control of endogenous tyramine 1 receptor (Tyr_1_R) expression (*Oct-TyrR-2A>TransTimer[sfGFP,RFP]*) and mean fluorescence intensity ratio values between sfGFP and RFP (Mean intensity (sfGFP/RFP)) in the central brain area of young (white) and aged (blue) brains, reflecting endogenous Tyr_1_R expression level. Data are presented as means ± SEM. Circles represent individual data points (mean values per brain). \*\**P* < 0.01 (Mann-Whitney *U*-test). **B-C)** Mean time-dependent changes in jGCaMP7f fluorescence intensity signal (Δ*F/F_0_*) before and after the addition of 10 µM octopamine (Oct; black lines) in Tdc2 neurons in brains of young (black) and aged (blue) *Drosophila* C) without (Control; *tdc2>jGCaMP7f;mCherry*) or D) with overexpression of Tyr_1_R (Tyr_1_R overexpression; *tdc2>jGCaMP7f;mCherry;Tyr_1_R*). Data are expressed in percentages as a relative change in the fluorescence (Δ*F/F_0_*), reflecting changes in cytosolic concentration of Ca^2+^. Numbers in parentheses depict the number of optic lobes analyzed. **D-E)** Mean maximum changes (Amplitude Δ*F/F_0_* (%); left), cumulative changes (Cumulative Δ*F/F_0_*; middle) and rate of changes in the jGCaMP7f Δ*F/F_0_* signal (Rate Δ*F/F_0_* (%/min); right) after the addition of 10 µM Oct in octopaminergic neurons in brains of young (white) and aged (blue) *Drosophila* E) without (Control) or F) with overexpression of the Tyr_1_R (Tyr_1_R overexpression). Amplitude and cumulative data are calculated within 3 min after stimulation. Data are presented as means ± SEM. Circles represent individual data points (value per optic lobe). \*\**P* < 0.01 (Mann-Whitney *U*-test)**F)** Mean climbing pass rate of aged *Drosophila* without (*tdc2/+* and *Tyr_1_R/+*) or with overexpression of the Tyr_1_R (*tdc2>Tyr_1_R*) in Tdc2 neurons. N=number of independent vials (experiments), n=number of consecutive recordings per vial. Data are normalized to mean climbing pass rate of flies overexpressing Tyr_1_Rs and presented as means ± SEM. Circles represent individual data points (mean values per vial (independent experiment)). \**P* < 0.05, \*\*\**P* < 0.001 (pairwise comparison, one-way ANOVA on ranks, followed by Dunn’s test).

### Restoration of tyramine 1 receptor expression reverses Ca^2+^ signalling and locomotion defects in aged *Drosophila* and prolongs lifespan

To test whether age-related decline in Tyr_1_R expression is causally related to aberrant Ca^2+^ signaling, we expressed the Ca^2+^ sensor jGCaMP7f selectively in (nor)adrenergic-like Tdc2-positive neurons (*tdc2>jGCaMP7f,mCherry*; Control). Ca²⁺ signals in optic lobe projections of Tdc2 neurons were measured during bath application of Oct. Indeed, Oct-induced Ca^2+^ transients were reduced 2-fold in Tdc2+ projections in brains from aged animals as compared to young ones (Control; Fig. 7B-D; Suppl. Table S10). This finding was in line with our observation of the general neuronal population (see Fig. 6).

Based on these findings, we hypothesized that selective overexpression of Tyr_1_R has the potential to reverse age-related phenotypes. Strikingly, when Tyr_1_Rs were overexpressed in Tdc2+ cells (*tdc2>Oct-TyrR;jGCaMP7f,mCherry*), Oct sufficed to induce Ca^2+^ transients with a magnitude comparable to those of young brains (Fig. 7C and E, Suppl. Fig. S11, Suppl. Table S10). Thus, reinstating Tyr_1_R expression in Tdc2 neurons in aged brains rescues impaired Ca^2+^ signalling.

Tdc2 neurons are known to play an important role in the regulation of locomotor function in *Drosophila* ^41^. Hence, we tested whether the age-related decline in climbing capability (Fig. 1) can also be restored by overexpression of Tyr_1_Rs in Tdc2+ cells (*tdc2>Tyr_1_R*). We compared the locomotor faculty of aged *tdc2>Tyr_1_R* with corresponding aged control groups (*tdc2/+* and *Tyr_1_R/+*) using an established climbing assay (^29^; Fig. 7F). The climbing pass rate of aged Tyr_1_R-overexpressing animals was nearly 2- and 3-fold higher than in respective control animals (Fig. 7F; Suppl. Table S11; Suppl. Fig. S12). These findings demonstrate the importance of Tyr_1_R-dependent Ca²⁺ signalling in Tdc2 neurons in maintaining motor function in old age.

This finding begged the question whether the animals’ lifespan could be prolonged by overexpression of Tyr_1_R. Thus, we compared the lifespan of *tdc2>Tyr_1_R* animals with corresponding control groups (*tdc2/+* and *Tyr_1_R/+*). Indeed, provision of Tyr_1_R in Tdc2+ cells prolonged lifespan by 23.3% (Mean lifespan *tdc2>Tyr_1_R*: 83.3 days, *tdc2/+*: 66.9 days, *Tyr_1_R/+:* 67.9 days, Suppl. Fig. S12).

Together, these findings show that restoring Tyr_1_R signalling in (nor)adrenergic-like Tdc2 neurons rescues octopamine-evoked Ca²⁺ responses, improves motor performance and extends overall lifespan.

## Discussion

### Neuronal D-glucose uptake declines with age, whereas neural L-lactate uptake is preserved

Neurons and glia are tightly coupled within the neural metabolic unit ^42^. Both cell types are capable of taking up D-glucose and L-lactate in the adult *Drosophila* brain ^43,44^ and have similar basal D-glucose and L-lactate levels (Fig. 2). Here we show that aging affects substrate metabolism differently in neurons and glia: Aging selectively reduced neuronal uptake of extracellular D-glucose, consistent with the decline in brain glucose utilization during aging and neurodegeneration ^2,9^. Accordingly, basal neuronal D-glucose levels were also reduced with age. This effect may be a reflection of faulty transport as several neuronal glucose transporters are known to be dysregulated in aged and neurodegenerative mammalian brains and in *Drosophila* ^9,45,46^. In contrast, L-lactate uptake remained stable in neurons and astrocytes, which implies that preserved lactate uptake may become important as neuronal glucose utilization declines.

### Old age affects octopamine-dependent metabolic responses selectively in astrocytes

Noradrenaline coordinates arousal, attention, stress responses, and cognition while increasing brain energy availability, including L-lactate delivery to active synapses. Age-related noradrenergic dysfunction ^20,22^ may therefore weaken neuron-astrocyte metabolic coupling and contribute to cognitive and motor decline. The *Drosophila* noradrenergic-like octopaminergic/tyraminergic system also changes with age and is associated with impaired motor function ^47^. Whether aging alters octopaminergic/tyraminergic modulation of the neuronal and/or astrocytic metabolism and thereby disrupts their metabolic coupling, remains unknown. Our cell-resolved measurements in living *Drosophila* brains revealed selective impairment of astrocytic responses to Oct, whereas key neuronal responses were preserved.

Neuronal free D-glucose levels did not change with Oct treatment or age, but Oct selectively increased astrocytic D-glucose uptake; this effect was markedly reduced in aged brains (Fig. 3). Oct application increased L-lactate levels in both cell types, but with a faster and larger response in neurons. Importantly, aging attenuated the astrocytic L-lactate response, while the neuronal response remained intact (Fig. 4). Thus, astrocytes appear to be the principal site of age-related decline in Oct-dependent metabolic responsiveness. This likely weakens neuromodulator-driven metabolic support of neuronal activity in aged *Drosophila* brains, as glial metabolic support was shown to be essential for neuronal survival ^48^. Given the high energetic demands and metabolic vulnerability of neurons ^9^, preservation of the Oct-driven neuronal L-lactate response may be a compensatory mechanism that helps sustain neuronal energy metabolism during aging.

Basal L-lactate levels did not differ between young astrocytes and neurons. Yet, MCT inhibition resulted in an approximately 4-fold higher rate of L-lactate accumulation in astrocytes compared to neurons (Suppl. Fig. S4). This indicates substantially greater intrinsic L-lactate production in astrocytes, which was also suggested in mammals ^11^. The Oct-evoked astrocytic L-lactate increase therefore likely reflects an increased rate in aerobic glycolysis, whereas the larger and faster neuronal response is more consistent with rapid uptake of extracellular L-lactate. In line with this interpretation, the astrocytes responded with a delay of more than 100 s after Oct-stimulation. Recently, an ANLS has been proposed in adult *Drosophila* brains during high-K⁺ stimulation (unpublished data Begić *et al.*, 2026) and in the larval brain during GABA receptor blockade with picrotoxin ^44^. Together, these findings support the model in which Oct stimulates astrocytic L-lactate production and its subsequent transfer to neurons. However, contribution of L-lactate through neuronal aerobic glycolysis ^49–51^ cannot be excluded.

### Aging reduces octopamine-mediated Ca^2+^ responses in both glia and neurons

Octopaminergic control of brain cell metabolism and locomotion likely involves activation of Oct/Tyr receptor pathways coupled to cAMP and/or Ca²⁺ signalling. We monitored these messengers in living brains to understand how aging could affect octopaminergic control of brain cell metabolism and locomotion.

Oct triggered a sustained, neuron-specific cytosolic cAMP increase that remained intact with age (Fig. 5). Similar Oct-evoked cAMP increases occur in mushroom body neurons ^52^. This finding is consistent with Gs-coupled Oct/Tyr receptor activation, including OctβRs ^32,53,54^, and may help preserve the neuronal L-lactate response during aging. A cAMP response was not detectable in astrocytes. Although an Oct-induced change in cAMP below or above the sensor’s detection threshold of 100 nM or 10 µM, respectively, cannot be excluded ^55^.

Oct evoked Ca²⁺ responses in both neurons and glia occur most likely through Gq-coupled receptor activation ^32,34^. Responses were faster and larger in neurons (Fig. 6; ^32^). Most strikingly, in aged brains Ca^2+^ signaling was almost lost in both neurons and astrocytes (Fig. 6). This finding aligns with the Ca²⁺ hypothesis of brain aging, which links disturbed Ca²⁺ regulation to neuronal dysfunction ^2,56^. Yet, change in Ca²⁺ signaling did not affect neuronal lactate responses, whereas astrocytic glucose and lactate responses declined. Neurons may preserve Oct-dependent metabolic responses through cAMP signalling despite age-related loss of Ca²⁺ signalling. Contrarily, astrocytes lack a detectable Oct-evoked cAMP response and could be more reliant on Ca²⁺-dependent metabolic regulation and consequently more vulnerable to its age-related decline.

The expression level of Tyr_1_R, a central nervous system Gq-coupled receptor (S. El-Kholy et al., 2015) linked to Ca²⁺ signaling ^34^, was significantly decreased in aged brains (Fig. 7). This is consistent with the age-related decline in octopamine and tyramine levels reported in *Drosophila* ^57^. Lower Tyr_1_R abundance would restrict receptor-mediated Ca²⁺ mobilization and weaken pathways required for neuronal and astrocytic function. Aging reduced neuronal Ca²⁺ mobilization after ionomycin and thapsigargin, indicating a deficit in intracellular Ca²⁺ availability. Reduced endoplasmic reticulum Ca²⁺ content may result from lower SERCA activity and/or impaired store-operated Ca²⁺ entry (SOCE). STIM and Orai activate SOCE after store depletion, allowing Ca²⁺ influx that refills the endoplasmic reticulum and sustains homeostasis ^58,59^. These defects could limit Ca²⁺ available for Oct/Tyr receptor signaling. In a *Drosophila* amyotrophic lateral sclerosis model, restoring the presynaptic voltage-gated Ca²⁺ channel *cacophony* (CaV2), essential for Ca²⁺ entry into presynaptic terminals and subsequent neurotransmitter release, rescues locomotor defects ^60^. However, we found no changes in cacophony expression levels ^61^ in young and aged brains (Suppl. Fig. S10), arguing against its involvement in physiological aging. Thus, the observed Ca²⁺ changes likely reflect impaired Tyr_1_R signalling combined with deficiencies in Ca²⁺ storage, refilling, receptor coupling, or a combination thereof.

### Tyramine 1 receptor overexpression in noradrenergic-like Tdc2 neurons restores Ca²⁺ signaling and locomotion in aged animals and prolongs lifespan

From humans to *Drosophila*, a hallmark feature of aging is the decline in locomotor ability. Here, we show that motoric decline in old age can be reversed by restoring Tyr_1_R expression in (nor)adrenergic-like Tdc2 neurons and thereby adequate Ca^2+^ signaling. Consitent with this finding, octopamine-dependent Ca²⁺ signaling has been shown to support metabolism and locomotion in *Drosophila* ^34,62,63^. Supporting the role of octopaminergic/tyraminergic Ca^2+^ signalling in motor control, OctαRs were also linked to Ca²⁺ signalling ^53^, regulate locomotion, as RNAi-mediated knockdown impaired climbing performance and induced muscle and leg morphological alterations ^62^. Moreover, the lifespan of animals overexpressing Tyr_1_R in Tdc2 neurons was prolonged, consistent with previous reports that increased neuronal synthesis of tyramine or octopamine in Tdc2 neurons, as well as dietary supplementation with these amines, can extend lifespan in *Drosophila* ^57^. In rodents, locomotion has been associated with Ca²⁺ signaling in neurons and astrocytes as well as engagement with the *locus coeruleus*– noradrenergic system ^64,65^. Thus, noradrenergic signalling may coordinate neural activity with metabolic status to accomodate the energetic demands of locomotion.

Many ligand–receptor systems exhibit autoregulatory negative-feedback mechanisms. Thus, reduced Tyr_1_R expression could result from decreased Oct and Tyr levels in aged *Drosophila* ^47,57^. In mammals, brain noradrenaline also declines with age ^66^, and ligand availability can modify neural adrenoceptor expression ^67^. Thus, other yet to be uncovered mechanisms underlie the observed downregulation of Tyr_1_R expression. Our findings show that the neuron-glial metabolic unit is reorganized during aging and collectively suggest a model in which octopamine-driven Ca^2+^ signaling enables adequate metabolic signaling that prevents the manifestation of age-releated symptoms.

In conclusion, aging selectively weakened Oct-dependent astrocytic metabolic response, yet preserved neuronal response, likey due to preserved Oct-mediated cAMP signals. On the other hand Oct-evoked Ca²⁺ signals declined in both neurons and glia, which associates with Tyr_1_R expression decline. Reinstaiting Tyr_1_R expression in octopaminergic Tdc2 neurons prolonged lifespan and rescues Ca²⁺ and locomotor deficits in aged animals, identifying a modifiable determinant of motor aging. Given the evolutionary conservation of neuromodulatory systems that regulate neural activity, metabolism, and behaviour, similar mechanisms may also operate in the mammalian brain during normal aging and neurodegenerative diseases, highlighting the translational relevance of these findings.

## Methods

Unless stated otherwise, all chemicals were of the highest quality and obtained from Sigma-Aldrich (Merck KGaA, Darmstadt, Germany).

### Fly maintenance and strains

Flies (*Drosophila melanogaster*) were raised at 25°C on standard cornmeal- and-molasses medium. Experiments and fly handling were performed in the laboratory for genetically modified organisms class 1 (GMO1; Permission No. 35417-2/2017-13; signed by Renata Perko).

The following fly strains were used (see also Supplemental Table S1 for detailed information): wild-type (*w1118*;;), *alrm-Gal4* (gift from Christian Klämbt;^68^), *elav-Gal4* (BDSC #8765;^69^), *R55B12-Gal4* (BDSC #39103;^70^), *R83E12-Gal4* (BDSC #40363;^70^), *R85G01-Gal4* (BDSC #40436;^70^), *SPG-Gal4* (BDSC #90883;^71^), *Oct-TyrR-2A-Gal4* (BDSC #86138), *tdc2-Gal4* (gift from Mareike Selcho;^72^), *UAS-jGCaMP7b* (BDSC #80907;^32,73^), *UAS-mCherry* (BDSC #52268), *UAS-Epac1-camps*^74^, *UAS-FLII^12^Pglu-700μδ6*^75^, *UAS-TransTimer[sfGFP,RFP]* (gift from Richard Binari;^40^), *UAS-Oct-TyrR* (BDSC #95289), and *UAS-jGCaMP7f* (BDSC #97031;^73^).

UAS-Laconic was generated as follows: The coding sequence of Laconic was amplified from pcDNA3.1 Laconic (Addgene #44238,^76^) using the primers caccCAGAATTCCACCACACTGGA and GCTGATCAGCGGTTTAAACTT and cloned into a pENTR-vector using pENTR™/D-TOPO® Cloning (Thermofisher). Then, the Laconic coding sequence was cloned via gateway cloning (Gateway LR, Invitrogen) into the vector pUASTattBrfa ^77^, which allows Φintegrase-mediated integration into the fly genome. The resulting vector has been integrated into the fly genome at landing site 86Fb.

Experiments were performed on brains isolated from young (up to 5 days old) and aged (≥30 days old) flies or age-matched living adult flies.

### Phalloidin staining and assessment of brain neurodegeneration

Brains of young and aged wild-type *Drosophila* flies (*w1118*) were dissected in ice-cold *Drosophila* Ringer’s solution containing (in mM): 3 CaCl_2_ × 2H_2_O, 182 KCl, 46 NaCl, 10 Tris Base (Carl Roth, Karlsruhe, Germany); pH was adjusted to 7.2 with 1 N HCl and the solution was filter sterilized (0.47 µm filter pores). Within 1 h of dissection, brains were fixed in 4% formaldehyde in phosphate buffered saline (PBS) for 45 min at RT, rinsed twice and washed 3×15 min in 1% PBT (PBS supplemented with 1% Triton X-100) while shaking gently. The brains were then placed in a blocking solution (1% PBT supplemented with 5% natural goat serum (NGS)) for 24 h at RT. Next, Alexa Fluor™ 647 dye conjugated with Phalloidin (1:20; Thermo Fisher Scientific, Cat. no. A22287) diluted in 1% PBT with 5% NGS was applied, and the brains were shaken for 24 h at 4°C. Finally, the brains were rinsed twice and washed 3×20 min in PBS before being mounted anterior side up onto glass slides using Vectashield mounting medium with DAPI (Vector Laboratories, United States).

Whole-brain imaging was performed using a confocal microscope Leica SP8 (Leica, Wetzlar, Germany) and 20×/0.75 IMM HC PL APO CS2 oil objective. Samples were exposed to 405 nm diode and 633 nm laser excitation and images at a resolution of 2048×2048 with a Z step of 1 µm were acquired. Emission was collected simultaneously with a HyD detector and band-pass emission filters of 410–550 nm (DAPI) and 638–750 nm (Alexa Fluor™ 647). Assessment of pathological vacuoles, which were shown to be associated with neurodegeneration in the *Drosophila* brain, was performed on blinded brain samples of young and aged flies using Fiji according to a previously published protocol ^27^.

### Climbing assay

Locomotor activity of young and aged wild-type flies (*w1118*) and of flies overexpressing the Tyr_1_R in octopaminergic neurons (Tdc2 neurons; *tdc2-Gal4>UAS-Tyr_1_R*), general neuronal population (*elav-Gal4*> *UAS-Tyr_1_R*), astrocytes (*alrm-Gal4*> *UAS-Tyr_1_R*) or astrocytes and neurons (*alrm;elav-Gal4>UAS-Tyr_1_R*) and their corresponding control strains (*tdc2-Gal4* (*tdc2/+*), *elav-Gal4* (*elav/+*), *alrm-Gal4* (*alrm/+*), *alrm;elav-Gal4* (*alrm;elav/+*) and *UAS-Oct-TyrR* (*Tyr_1_R/+*)) was assessed at room temperature (RT) using a climbing assay method ^28,29^. Prior to the assay, flies of each genotype were transferred into empty vials (∼10 animals/vial). Climbing capability was monitored for 10 s over 10 consecutive recordings/vial. ∼ 100 animals were recorded for each experimental group. For each recording, the number of flies that climbed above an 8-cm mark in height after being tapped to the bottom of the vial was counted. The mean number of flies that surpassed the 8-cm mark across all 10 recordings per vial was calculated, representing one independent experiment (datapoint). Climbing performance was expressed as the mean climbing pass rate (percentage of flies that surpassed the 8-cm mark).

### Whole brain (*ex vivo*) imaging of second messengers and metabolites

#### Experimental solutions

Imaging of cytosolic Ca^2+^ and cAMP dynamics was performed in an experimental solution composed of (in mM): 113.5 NaCl (Carlo Erba 479687), 5.4 KCl, 1.2 CaCl_2_, 0.8 MgCl_2_, 10 HEPES, and 3 D-glucose; pH was adjusted to 7.2 with NaOH. Imaging of cytosolic D-glucose and L-lactate dynamics was performed in an experimental solution composed of (in mM): 113.5 NaCl, 5.4 KCl, 1.2 CaCl_2_, 0.8 MgCl_2_, 10 HEPES, 3 D-glucose, 50 sucrose, and 5 trehalose dihydrate, pH was adjusted to 7.2 with NaOH. All solutions were filter sterilized (0.47 µm filter pores). Osmolality was measured with Osmomat 030 (Gonotech GmbH, Germany).

#### Brain dissection

Brain dissection was performed in ice-cold Ca^2+^-free experimental solutions; NaCl was used to compensate for the osmolality. After dissection, brains were transferred to coverslips precoated with 0.1% poly-L-lysine with the anterior side facing down, which were previously mounted onto a recording chamber. Brains were supplied with experimental solution and incubated for 15 min at RT before the experiments ^32^.

#### Fluorescence measurements of cytosolic Ca^2+^ dynamics

Brains of young and aged *Drosophila* co-expressing Ca^2+^ sensor jGCaMP7b and mCherry in astrocytes (*alrm>jGCaMP7b;mCherry*), neurons (*elav>jGCaMP7b;mCherry*) and cortex glia (*R55B12>jGCaMP7b;mCherry*) were imaged with a Zeiss LSM 800 confocal microscope (Zeiss, Jena, Germany) using a Plan-Apochromat 40×/1.3 Oil objective and 488 nm (Ca^2+^ sensor) and 561 nm (mCherry) laser excitation. Images were acquired every 1 s (neurons) or 3 s (neurons and neuroglia), total recording time was 900 s. At 300 s, brains were exposed by bath application to 10 μM octopamine (Oct). In some experiments, brains co-expressing jGCaMP7b and mCherry in neurons (*elav>jGCaMP7b;mCherry*) were bath-stimulated with 1 μM thapsigargin (TG), an inhibitor of the SERCA pump that prevents the reuptake of Ca^2+^ into the ER ^38^ or 10 μM ionomycin (Iono), a Ca^2+^ ionophore that facilitates the transport of Ca^2+^ across cellular membranes^39^. In a separate set of experiments, using a Plan-Apochromat 20×/0.8 objective, brains co-expressing Ca^2+^ sensor jGCaMP7f and mCherry, either without or with overexpression of Tyr_1_Rs (Oct-TyrRs) in octopaminergic neurons (*tdc2>jGCaMP7f,mCherry* or *tdc2>Oct-TyrR;jGCaMP7f,mCherry*, respectively), were imaged before (300 s baseline recording) and after bath-stimulation (600 s) with 10 μM Oct. In control experiments, brains were stimulated by bath application of experimental solution only (vehicle; Control).

Fluorescence emissions of jGCaMP7b/jGCaMP7f and mCherry were collected simultaneously using 410–546 nm and 640–700 nm band-pass emission filters, respectively. To quantify the jGCaMP7b/jGCaMP7f and mCherry fluorescence signal intensities, a region of interest (ROI) was selected for randomly chosen single cell somata or single brain regions (optic lobe; experiments on octopaminergic neurons) using Zeiss Zen Imaging or Fiji software. The mean fluorescence intensity of all pixels within each ROI was extracted for each time point, followed by background fluorescence subtraction from the fluorescence signal of the single cell soma or optic lobe. Changes in jGCaMP7b, jGCaMP7f or mCherry signal are expressed as a relative change in fluorescence, Δ*F/F_0_ = (F-F_0_)/F_0_*, where *F_0_* denotes the signal intensity in the first 300 s of recordings (baseline). An increase in the jGCaMP7b/jGCaMP7f signal reflects an increase in the cytosolic concentration of Ca^2+^. The mCherry signal served as a reference for the jGCaMP7b and jGCaMP7f signals, as it is not sensitive to changes in cytosolic Ca^2+^ ^32^.

The cumulative change in the Δ*F/F_0_* signal was calculated as the sum of all Δ*F/F_0_* values recorded at each time point over a period of 600-s or 180-s (octopaminergic neurons) after applying stimulus. The rate of the Δ*F/F_0_* signal change was determined as the slope of the linear regression function [Δ*F/F_0_* (%) = slope (%/min) × time (min)] fitting the initial signal change. The maximum change (amplitude) in the Δ*F/F_0_* jGCaMP7f signal in octopaminergic neurons was determined by subtracting the baseline mean Δ*F/F_0_* signal value (first 300 s) from the maximum Δ*F/F_0_* signal value observed within the first 3 min after stimulation (peak value ± 1-2 data points). All acquired Δ*F/F_0_* signals (responsive and unresponsive) were included in the analysis.

#### FRET measurements of cytosolic cAMP, D-glucose and L-lactate dynamics

Brains isolated from young and aged *Drosophila* expressing cAMP (Epac1-camps), D-glucose (FLII^12^Pglu-700μδ6), and L-lactate (Laconic) FRET-based sensors in astrocytes (*alrm>Epac1-camps*, *alrm>FLII^12^Pglu-700μδ6* and *alrm>Laconic*, respectively), neurons (*elav>Epac1-camps*, *elav>FLII^12^Pglu-700μδ6* and *elav>Laconic*, respectively) and cortex glia (*R55B12>Epac1-camps*, *R55B12>FLII^12^Pglu-700μδ6* and *R55B12>Laconic*, respectively) were imaged with a Zeiss LSM 780 confocal microscope (Zeiss, Jena, Germany) using a Plan-Apochromatic 40×/1.3 Oil objective and 458 nm laser excitation. Images were acquired every 3 s (cAMP) or 10 s (D-glucose, L-lactate), total recording time was 900 s. At 300 s, the brains were exposed to bath application of 10 µM Oct, 20 mM D-glucose or 30 mM L-lactate. In addition, brains isolated from young *Drosophila* expressing the L-lactate sensor Laconic in neuropil ensheathing glia (*R83E12>Laconic*), perineurial glia (*R85G01>Laconic*) and subperineurial glia (*SPG>Laconic*) were after 300 s stimulated by bath application to 10 µM Oct. In control experiments, brains were bath-stimulated with experimental solution only (vehicle; Control). Two sets of experiments were performed using α-cyano-4-hydroxycinnamate (CHC). In the first, brains expressing Laconic in astrocytes or neurons, were after 300 s of baseline bath-stimulated with 6 mM CHC and recorded for another 25 min (30 min total). In the second, brains were preincubated for 15 min at room temperature in experimental solution containing either 6 mM CHC or CHC vehicle. After 300 s of baseline recording, brains were bath-stimulated with 10 µM Oct for 600 s, followed by 30 mM L-lactate for 600 s (1,500 s total) while maintaining Oct concentration, in the presence or absence of CHC.

Fluorescence emission of CFP and YFP variants in the FRET sensors was collected simultaneously using 473-499 nm (CFP, ECFP, mTFP) and 508-534 nm (YFP, EYFP, Venus) band-pass emission filters. For each fluorescent protein, the mean fluorescence intensity was measured at each time point in randomly selected single cell somata (astrocytes, neurons and cortex glia) or single or multiple brain areas (neuropil ensheathing glia, perineurial and subperineurial glia) using the ROI tool in Zeiss Zen Imaging software. The background fluorescence for each protein was subtracted from the individual cell/region fluorescence signal. The sensor fluorescence ratio signals were calculated, CFP/YFP (Epac1-camps), EYFP/ECFP (FLII^12^Pglu-700μδ6) and mTFP/Venus (Laconic), reflecting real-time dynamics in cytosolic cAMP, D-glucose and L-lactate, respectively. Fluorescence ratio signals were normalized to 1 and expressed as the percentage change relative to the baseline ratio signal. The mean ratio signal change was determined by subtracting the mean ratio signal value of 100 s before stimulation from the mean ratio signal value of the last 100 s upon stimulation. The cumulative change in the ratio signal was calculated as the sum of all ratio signal values recorded at each time point over a period of 600-s after applying stimulus. The initial rate of ratio signal change (CFP/YFP, EYFP/ECFP or mTFP/Venus) was calculated as the slope of the linear regression function [ratio signal (%) = slope (%/min) × t (min)] fitting the initial ratio signal change.

For calculation of the basal Laconic and FLII^12^Pglu-700μδ6 ratio signals, absolute basal sensor signals, reflecting basal L-lactate and D-glucose levels, respectively, were normalized to the corresponding saturated sensor signals, following the approach described by Mächler et al. ^78^. We assumed that 30 mM L-lactate and 20 mM D-glucose, concentrations exceeding the sensors’ detection ranges, fully saturated the sensors. The reported detection ranges are 1 µM– 10 mM for Laconic L-lactate ^79^ and 50 µM–11 mM for FLII^12^Pglu-700µδ6 D-glucose sensing ^80^. Because we could not account for potential cell volume changes or intrinsic metabolism during the measurements, some degree of misinterpretation cannot be entirely excluded ^81^.

### Seahorse measurements

Seahorse XFe96 analyser (Agilent, Santa Clara, USA) was set to 25°C with the heat on and placed in a temperature-controlled room set to 4°C. Brains of young and aged wild-type *Drosophila* (*w1118*) were dissected in ice-cold isolating solution containing 3 mM CaCl_2_ × 2H₂O, 182 mM KCl, 46 mM NaCl, 10 mM Tris base (pH = 7.2) and transferred to 96-well metabolic assay cell plates (Agilent) containing XFe96 assay media supplemented with 10 mM glucose and 10 mM sodium pyruvate (pH = 7.4). Brains were carefully centred between the three raised spheres and lowered to the bottom of the wells. Then, micro-tissue restraints (manufactured by Arizona State University, Instrument Design and Fabrication Core, Tempe, USA; patent no. 62/491,431, M. Tipping) were placed inside each well to keep the brains in place during the experiment. The starting volume of the XFe96 assay media was 180 µL.

Within 30 min of brain dissection, the metabolic profile of young and aged brains under basal and Oct-stimulated conditions was evaluated using the Mito Stress Test Kit (Agilent, Santa Clara, USA) in accordance with the manufacturer’s instructions and previously published protocols ^82,83^ with minor adjustments. Oxygen consumption rates (OCR) and extracellular acidification rates (ECAR) were measured over 8 cycles for each experimental condition. Each cycle consisted of one minute mixing, no waiting time, and three minutes of measuring. The experimental condition included: basal, Oct (1 mM) and oligomycin (10 µM) and rotenone/antimycin A (5 µM) in the presence of 1 mM Oct (Oligo+Oct and Rot/AA+Oct, respectively). Oct, Oligo+Oct and Rot/AA+Oct were injected at the 8^th^, 16^th^ and 24^th^ cycle, respectively. The total measurement time was ∼150 min. Oligo and Rot/AA are mitochondrial inhibitors used to evaluate mitochondrial respiratory chain function and metabolic pathways. To exclude potential effects of micro-tissue restraints on data acquisition, six wells per experiment contained only assay media and the restraints.

The metabolic profiles of young and aged *Drosophila* brains were assessed by calculating the ratio between OCR and ECAR (OCR/ECAR ratio). For each cycle, the mean OCR/ECAR ratio acquired from all brains within the experimental age group (typically in triplicate; 3 brains from young and 3 brains from aged *Drosophila*) was calculated, representing an independent experiment. Each dataset for the young and aged group was acquired from 5 independent experiments (i.e., 5 different runs on Seahorse analyzer).

### Evaluation of brain endogenous tyramine 1 receptor expression level

Brains of young and aged *Drosophila* expressing UAS-TransTimer, i.e., a fast-folding GFP (sfGFP) and a slow-folding RFP under the control of endogenous *Tyr_1_R* promotor (*Oct-TyrR-2A-Gal4>UAS-TransTimer[sfGFP,RFP]*) were dissected in ice-cold *Drosophila* Ringer’s solution. Immediately after dissection, brains were fixed in 4% formaldehyde in PBS for 30 min at RT, then rinsed twice and washed 3×15 min in PBS while shaking gently. After incubating in Vectashield mounting medium overnight at 4°C, the brains were mounted anterior side up onto glass slides using Vectashield mounting medium with DAPI. Whole-brain imaging was performed using a confocal microscope Leica SP8 (Leica, Wetzlar, Germany) and 20×/0.75 IMM HC PL APO CS2 oil objective. Brains were exposed to 488 nm (sfGFP) or 561 nm (RFP) laser excitation and images at a resolution of 1024×1024 with a Z-step of 2 µm were acquired. Emission was collected with a HyD detector and band-pass emission filters of 495–540 nm (sfGFP) and 580–650 nm (RFP). The analysis was carried out using Fiji and Excel on maximum intensity projection images for each brain separately. We determined the total sum of intensity units (RawIntDen) for sfGFP and RFP. Analysis was performed in the central brain region because in other brain regions autofluorescence was observed in the RFP channel. The Tyr_1_R expression level is presented as the ratio between sfGFP and RFP fluorescence signal intensity for young and aged brains separately.

### Evaluation of Cacophony expression levels

Brains of young and aged *Cac^sfGFP-N^*flies were dissected, mounted and imaged as described above. Fly brains were imaged using a 488 nm excitation wavelength, with fluorescence emission collected between 495 and 550 nm. Analysis was performed using ImageJ on maximum intensity projection images for each brain separately. The total sum of intensity units (RawIntDen) was determined for sfGFP on antennal lobes.

#### Protein extraction and immunoblotting

Adult wildtype (3 and 35 days) and *Cac^sfGFP-N^*(3 and 35 days) flies were used to compare protein abundance in young and aged animals. For each sample, 30 fly heads were collected, snap-frozen in liquid nitrogen and stored at −80 °C. Heads were homogenized on ice in 60 µl 2% SDS containing protease inhibitor (1:1,000), followed by addition of 6 µl 10% Triton X-100 and 0.6 µl 100 mM DTT. Samples were incubated on ice for 10 min with vortexing after 5 min, followed by centrifugation at 14,000 rpm for 30 min and subsequently 15 min at 4 °C. The supernatant was collected after each centrifugation.

Protein concentration was determined using the DC Protein Assay (Bio-Rad) with BSA standards. Samples were diluted 1:10 in lysis buffer, and absorbance was measured at 750 nm using a NanoPhotometer NP80 (Implen). Equal amounts of protein (100 or 200 µg) were separated on 4–12% Tris-glycine gels (Invitrogen) at 130 V for 75 min and transferred to nitrocellulose membranes using an iBlot 2 system (Invitrogen; 15 V, 7 min). Membranes were stained with Ponceau S for 5 min to assess protein transfer.

Following protein transfer, membranes were blocked for 1 h at room temperature in a 1:1 mixture of 1× PBS and Intercept Blocking Buffer (PBS; LI-COR). Membranes were incubated overnight at 4 °C with rabbit anti-GFP antibody (1:500; Invitrogen, G10362) and mouse anti-β-tubulin antibody (1:2,000; DSHB, E7c). After two rinses and one 5-min wash with 0.1% PBST, membranes were incubated for 1 h at room temperature with IRDye 800CW goat anti-rabbit (1:15,000; LI-COR) and IRDye 680RD goat anti-mouse (1:15,000; LI-COR) secondary antibodies. Membranes were washed as above and imaged using a LI-COR imaging system.

#### Western blot analysis

Cac^sfGFP-N^ expression was quantified by densitometric analysis of western blot bands using ImageJ. Band intensities were measured using the Gel Analysis tool, with an equivalent region defined for each band. Background signal was measured from an adjacent area of the membrane and subtracted from each measurement. Cac^sfGFP-N^band intensities were normalized to the corresponding tubulin signal and expressed relative to the respective control group.

### Lifespan Assay

For lifespan assay, 10 adult flies per genotype (5 females and 5 males) were collected and maintained at 25 °C under a 12-h light/12-h dark cycle. Flies were transferred to fresh vials every 2 days and survival was assessed daily.

### Statistical analysis

We selected the parametric test based on a normality analysis performed manually in GraphPad Prism (GraphPad Software, Inc., USA) software. Statistical significance between two or more groups was determined by Mann-Whitney *U*-test or one-way ANOVA on ranks, followed by Dunn’s test, respectively, unless stated otherwise. *P* < 0.05 was considered statistically significant. Whole brain (ex vivo) real-time experimental data were obtained from multiple cells or from multiple or one larger region (e.g. optic lobe) within each brain. The mean value of all analyzed cells in an individual brain, or of region(s), represents an independent experiment. The number of independent experiments (i.e., brains or regions) and the number of analyzed cells are provided in the figure legends and supplemental material. Data are presented as means ± standard error of the mean (SEM).

## Acknowledgements

Stocks obtained from the Bloomington Drosophila Stock Center (NIH P40OD018537) were used in this study. We thank Primož Runovc, Erika Rus, Miha Pate, Paula Beckmann and Lena Abicht for their technical assistance.

## Abbreviations

ANLS: astrocyte–neuron L-lactate shuttle
BDSC: Bloomington Drosophila Stock Center
BSA: bovine serum albumin
cAMP: cyclic adenosine monophosphate
CHC: α-cyano-4-hydroxycinnamate
CFP: cyan fluorescent protein
DAPI: 4′,6-diamidino-2-phenylindole
DTT: dithiothreitol
ECAR: extracellular acidification rate
ECFP: enhanced cyan fluorescent protein
ER: endoplasmic reticulum
EYFP: enhanced yellow fluorescent protein
FRET: Förster resonance energy transfer
GFP: green fluorescent protein
GMO: genetically modified organism
HEPES: 4-(2-hydroxyethyl)-1-piperazineethanesulfonic acid
Iono: ionomycin
MCT: monocarboxylate transporter
mTFP: monomeric teal fluorescent protein
NGS: normal goat serum
Oct: octopamine
OCR: oxygen consumption rate
OctR: octopamine receptor
Oct-TyrR: octopamine/tyramine receptor
PBS: phosphate-buffered saline
PBT: phosphate-buffered saline containing Triton X-100
PBST: phosphate-buffered saline containing Tween 20
RFP: red fluorescent protein
RNAi: RNA interference
ROI: region of interest
RT: room temperature
SDS: sodium dodecyl sulfate
SEM: standard error of the mean
SERCA: sarco/endoplasmic reticulum Ca²⁺-ATPase
sfGFP: superfolder/fast-folding green fluorescent protein
SOCE: store-operated Ca²⁺ entry
SPG: subperineurial glia
Tdc2: tyrosine decarboxylase 2
TG: thapsigargin
Tyr: tyramine
Tyr_1_R: tyramine 1 receptor
UAS: upstream activating sequence
YFP: yellow fluorescent protein

