## Supplemental Materials for "Restoration of (nor)adrenergic-like Ca²⁺ signalling counteracts age-related motor decline"

Running title:

(Nor)adrenergic-like $Ca^{2+}$ Signaling reverses Age-Related Motor Decline

Anemari Horvat[1,2], Urška Černe[1], Anne-Kristin Dahse[3], Lara-Sophie Brodmerkel[3], Aljaž Bratanič[1], Madlen Matz-Soja[4], Stefanie Schirmeier[5], Robert Zorec[1,2], Nicole Scholz[3,*], Nina Vardjan[1,2,*]

[1]*Laboratory of Neuroendocrinology-Molecular Cell Physiology, Institute of Pathophysiology, Faculty of Medicine, University of Ljubljana, Slovenia*
[2]*Laboratory of Cell Engineering, Celica Biomedical, Slovenia*
[3]*Division of General Biochemistry, Rudolf Schönheimer Institute of Biochemistry, Medical Faculty, Leipzig University, Germany*
[4]*Research Laboratory for Clinical and Experimental Hepatology, Central Research Building, Leipzig University, Germany*
[5]*Faculty of Biology, Technische Universität Dresden, Dresden, Germany*

**Funding:** This work was supported by Slovenian Research and Innovation Agency (P3-0310, J3-50104, N3 0470, J4-60077, I0-0034, I0-0022, I0-0048 (CIPKeBiP)), COST Actions CA18133 (ERNEST) and CA25102, and the Deutsche Forschungsgemeinschaft to N.S. (265903901, FOR 2149/P01 [SCHO1791/1-2] and CRC 1423 project number 421152132, subproject B06) and to S.S. (468211919).

**Acknowledgements:** Stocks obtained from the Bloomington Drosophila Stock Center (NIH P40OD018537) were used in this study. We thank Primož Runovc, Erika Rus, Miha Pate, Paula Beckmann and Lena Abicht for their technical assistance.

#### Supplemental Figures and Figure Legends

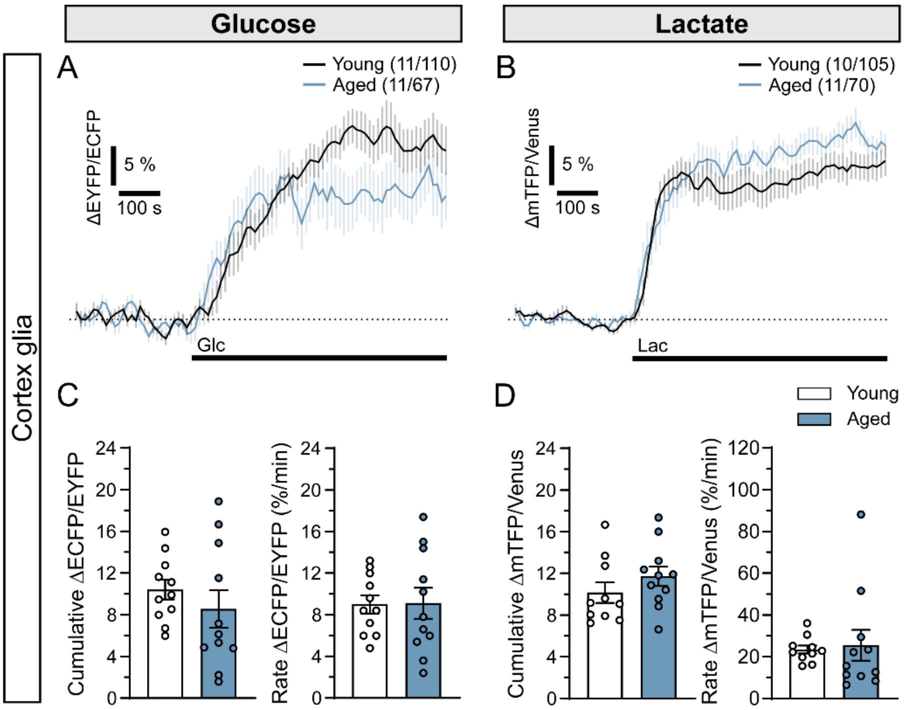


**Supplemental Fig. S1: Cytosolic D-glucose and L-lactate levels in cortex glia upon exposure of brains of young and aged *Drosophila* to elevated extracellular D-glucose and L-lactate.**

**A-B)** Mean time-dependent changes in the A) FLII$^{12}$Pglu-700µδ6 and B) Laconic fluorescence ratio signal (ΔEYFP/ECFP and ΔmTFP/Venus, respectively) before and after the addition of A) 20 mM D-glucose (Glc; black line) and B) 30 mM L-lactate (Lac; black line) in cortex glia in brains of young and aged *Drosophila*. Data are expressed in percentages as a change in fluorescence ratio signal (ΔEYFP/ECFP and ΔmTFP/Venus) relative to the baseline values, reflecting changes in cytosolic D-glucose and L-lactate levels. Numbers in parentheses depict the number of brains and cells analyzed. **C-D)** Mean cumulative changes (Cumulative ΔEYFP/ECFP and Cumulative ΔmTFP/Venus; left) and rate of changes in the C) FLII$^{12}$Pglu-700µδ6 and D) Laconic ratio signal (Rate ΔEYFP/ECFP (%/min) and Rate ΔmTFP/Venus; right) after the addition of C) 20 mM Glc and D) 30 mM Lac in cortex glia in brains of young (white) and aged (blue) *Drosophila*. Data are presented as means ± SEM. Circles represent individual data points (mean values of all analyzed cortex glia within individual brain).

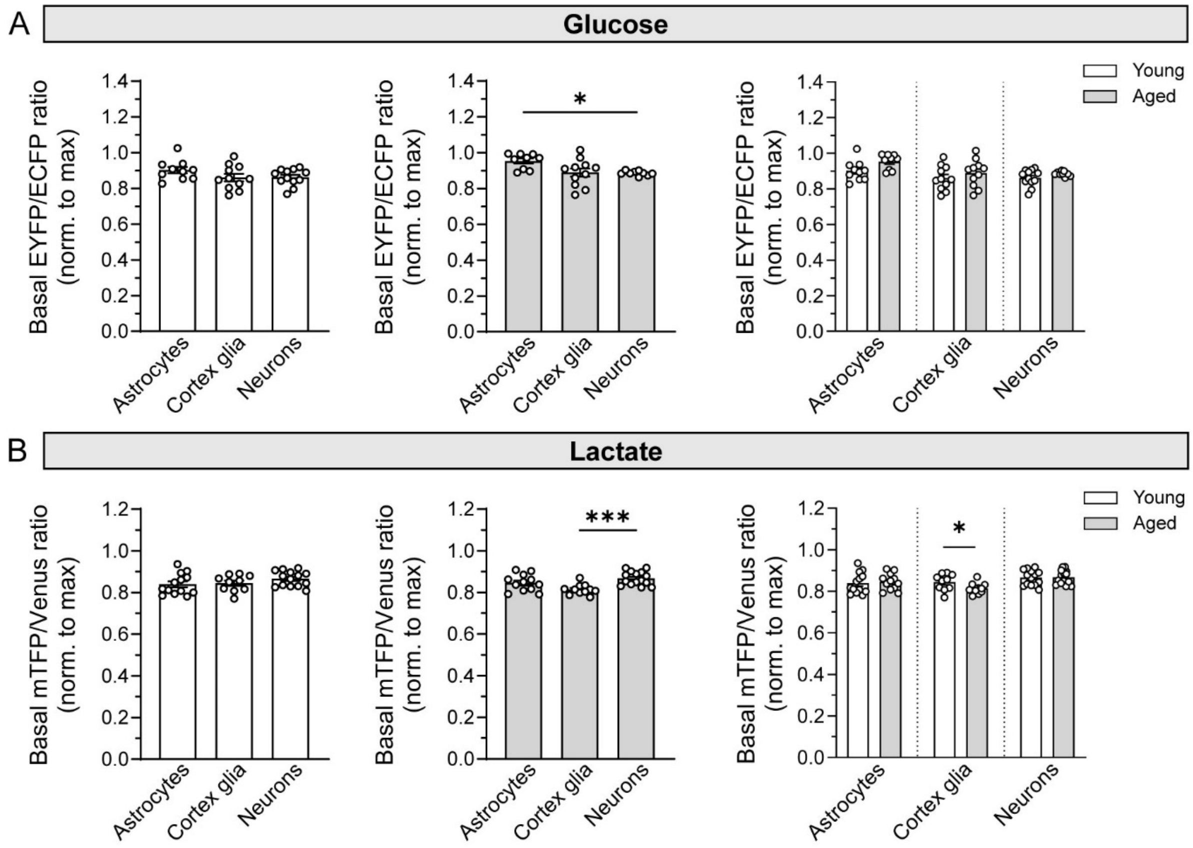


**Supplemental Fig. S2: Basal metabolite levels in glial cells and neurons in brains of young and aged *Drosophila*.**

**A-B)** Mean baseline values normalized to maximal response upon addition of 20 mM D-glucose and 30 mM L-lactate of A) FLII$^{12}$Pglu-700µδ6 (EYFP/ECFP) and B) Laconic (mTFP/Venus) fluorescence ratio signal, reflecting basal cytosolic D-glucose and L-lactate levels, respectively, in astrocytes, cortex glia and neurons in brains of young (white) and aged (grey) *Drosophila*. Data are presented as means ± SEM. White circles represent individual data points (mean values of all analyzed cells within individual brain/optic lobe). Left and middle graphs: $^{*}P < 0.05$, $^{***}P < 0.001$ (Brown-Forsythe and Welch ANOVA tests, followed by Dunnett's T3 test (B, middle panels) or One-way ANOVA on ranks, followed by Dunn's test (A, middle panels)); Right graphs: $*P < 0.05$ (Unpaired t-test).

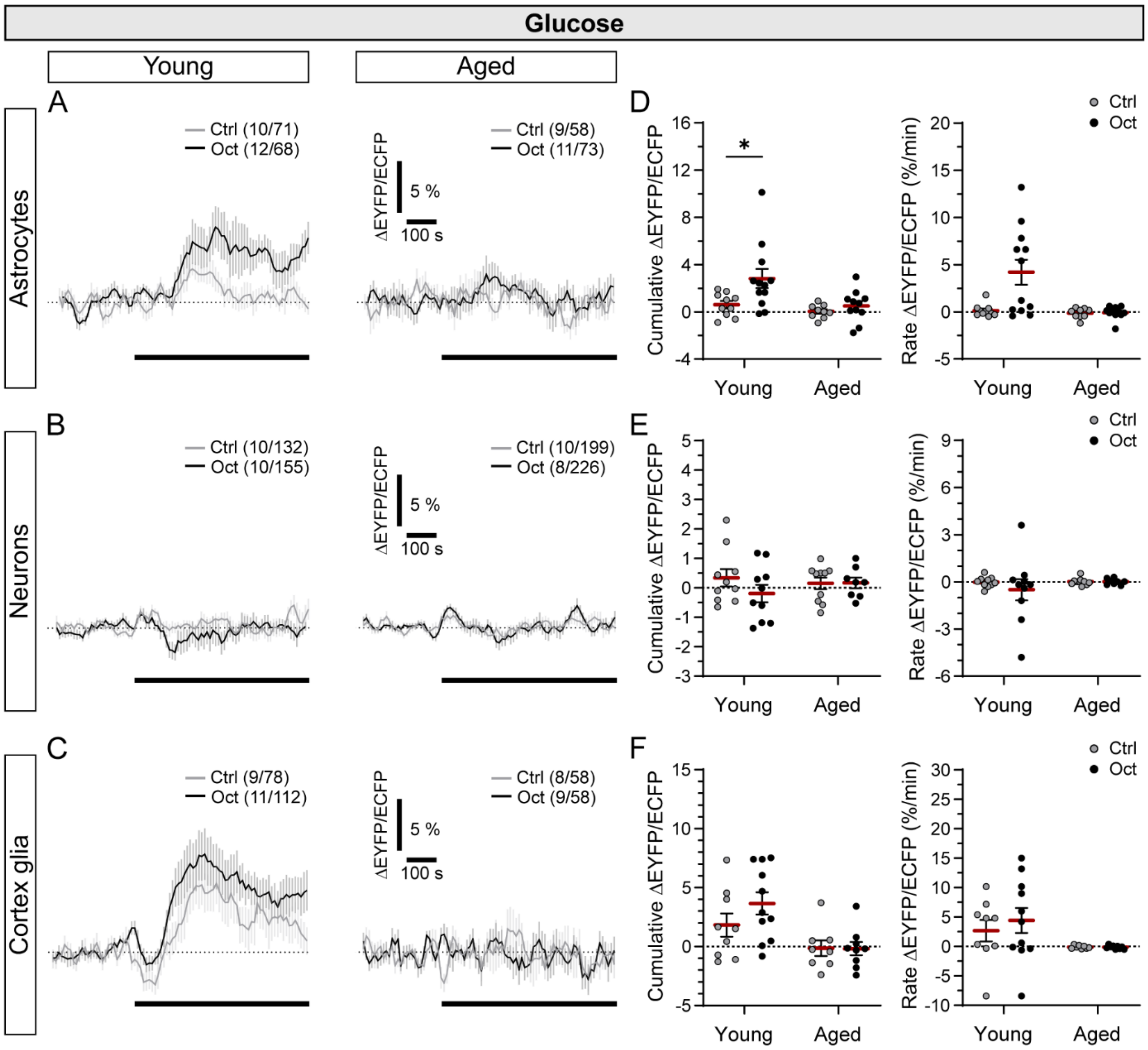


**Supplemental Fig. S3: Comparison of cytosolic D-glucose levels in glial cells and neurons in brains of young and aged *Drosophila* upon vehicle versus octopamine stimulation.**

**A-C)** Mean time-dependent changes in the FLII$^{12}$Pglu-700µδ6 fluorescence ratio signal (ΔEYFP/ECFP) before and after the addition of experimental solution (vehicle; Ctrl; grey) and 10 µM octopamine (Oct; black) in A) astrocytes (*alrm>FLII$^{12}$Pglu-700µδ6*), B) neurons (*elav>FLII$^{12}$Pglu-700µδ6*), and C) cortex glia (*R55B12>FLII$^{12}$Pglu-700µδ6*) in brains of young (left panels) and aged (right panels) *Drosophila*. Data are expressed in percentages as a change in fluorescence ratio signal (ΔEYFP/ECFP) relative to the baseline values, reflecting changes in cytosolic D-glucose levels. Numbers in parentheses depict the number of brains and cells analyzed. **D-F)** Scatter dot plots representing cumulative changes (Cumulative ΔEYFP/ECFP; left) and rate of changes in the FLII$^{12}$Pglu-700µδ6 ratio signal (Rate ΔEYFP/ECFP (%/min); right) after the addition of Ctrl (grey) and 10 µM Oct (black) in D) astrocytes, E) neurons and F) cortex glia in brains of young and aged *Drosophila*. Red lines represent mean of all data points; SEM is shown in black. Circles represent individual data points (mean values of all analyzed cells within individual brain/optic lobe). $*P < 0.05$ (Mann-Whitney *U*-test).

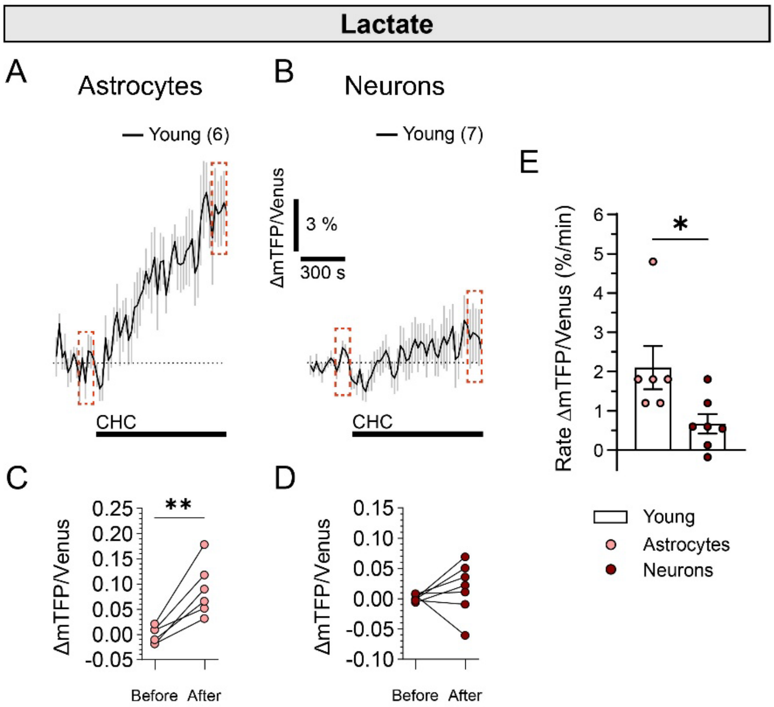


**Supplemental Fig. S4: In young brains, inhibition of monocarboxylate transporter activity elevates cytosolic lactate levels only in astrocytes**

**A-B)** Mean of time-dependent changes in the Laconic fluorescence ratio signal (ΔmTFP/Venus) before and after the addition of MCT inhibitor α-cyano-4-hydroxycinnamate (CHC; 6 mM) in A) astrocytes (*alrm>Laconic*) and B) neurons (*elav>Laconic*) in young *Drosophila* brains. Data are expressed in percentages as a change in fluorescence ratio signal (ΔmTFP/Venus) relative to baseline values. The mean vehicle (control) signal was subtracted from the CHC-induced signal for each cell type. Numbers in parentheses indicate the number of brains analyzed. **C-D**) Paired comparison of the mean ratio signals collected 100 s prior stimulation and the mean ratio signal of the last 100 s of 6 mM CHC stimulation in astrocytes and neurons in brains of young *Drosophila* (indicated by red dotted squares in A and B). $^{**}P < 0.01$ (Paired t-test). **E)** Scatter dot plot representing rate of changes in the Laconic ratio signal (Rate ΔmTFP/Venus (%/min)) after the addition of CHC in astrocytes and neurons in brains of young *Drosophila*. Circles represent individual data points (mean values of all analyzed cells within individual optic lobe). $^{*}P < 0.05$ (Mann-Whitney *U*-test).

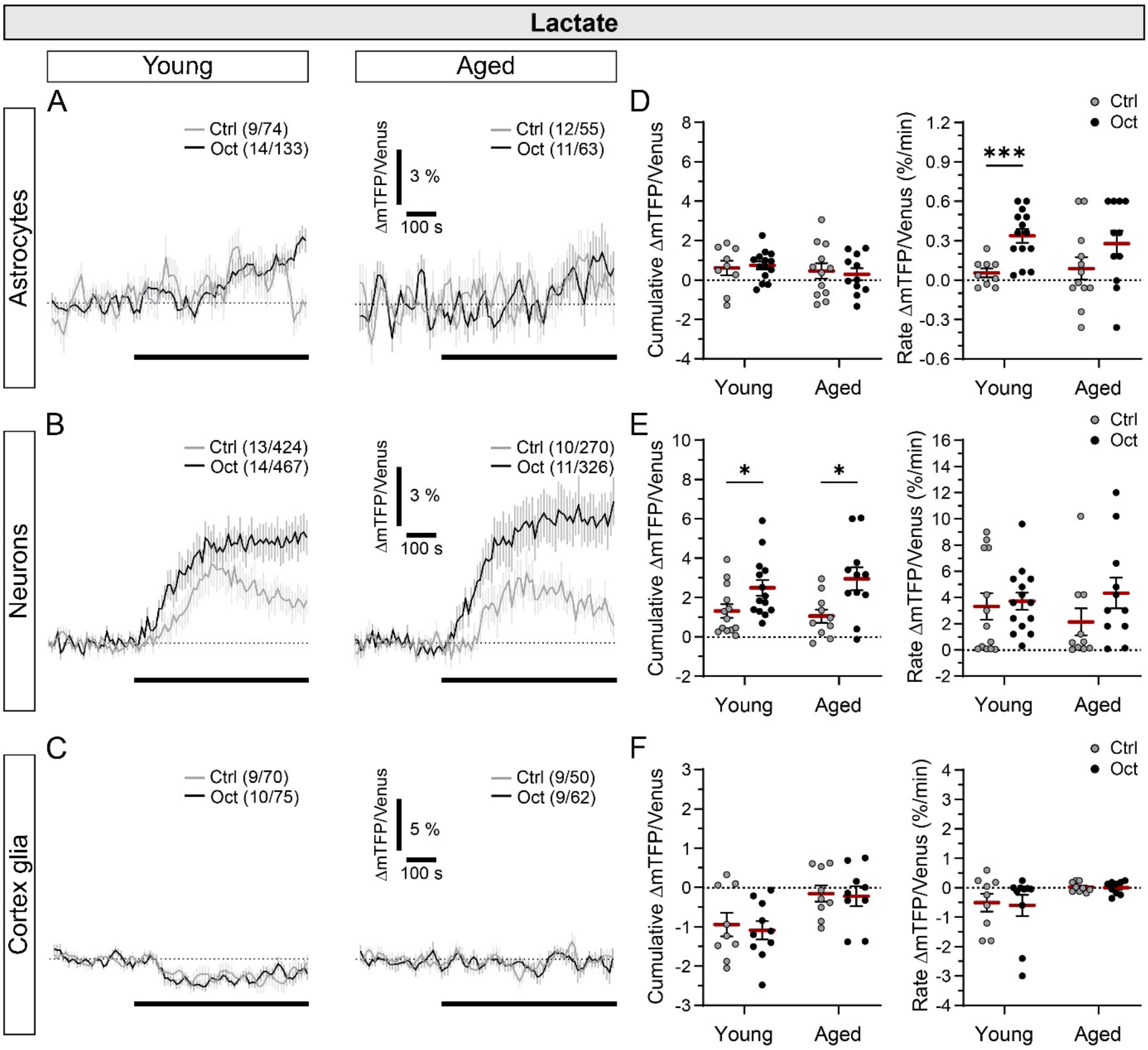


**Supplemental Fig. S5: Comparison of cytosolic L-lactate levels in glial cells and neurons in brains of young and aged *Drosophila* brains upon vehicle and octopamine stimulation. A-C)** Mean time-dependent changes in the Laconic fluorescence ratio signal (ΔmTFP/Venus) before and after the addition of experimental solution (vehicle; Ctrl; grey) and 10 µM octopamine (Oct; black) in A) astrocytes (*alrm>Laconic*), B) neurons (*elav>Laconic*), and C) cortex glia (*R55B12>Laconic*) in brains of young (left panels) and aged (right panels) *Drosophila*. Data are expressed in percentages as a change in fluorescence ratio signal (ΔmTFP/Venus) relative to the baseline values, reflecting changes in cytosolic L-lactate levels. Numbers in parentheses depict the number of brains and cells analyzed. **D-F)** Scatter dot plots representing cumulative changes (Cumulative ΔmTFP/Venus; left) and rate of changes in the Laconic ratio signal (Rate ΔmTFP/Venus (%/min); right) after the addition of Ctrl (grey) and 10 µM Oct (black) in D) astrocytes, E) neurons and F) cortex glia in brains of young and aged *Drosophila*. Red lines represent mean of all data points; SEM is shown in black. Circles represent individual data points (mean values of all analyzed cells within individual brain/optic lobe). $*P < 0.05$, $***P < 0.001$ (Unpaired t-test with Welch's correction (D) or Mann-Whitney *U*-test (E)).

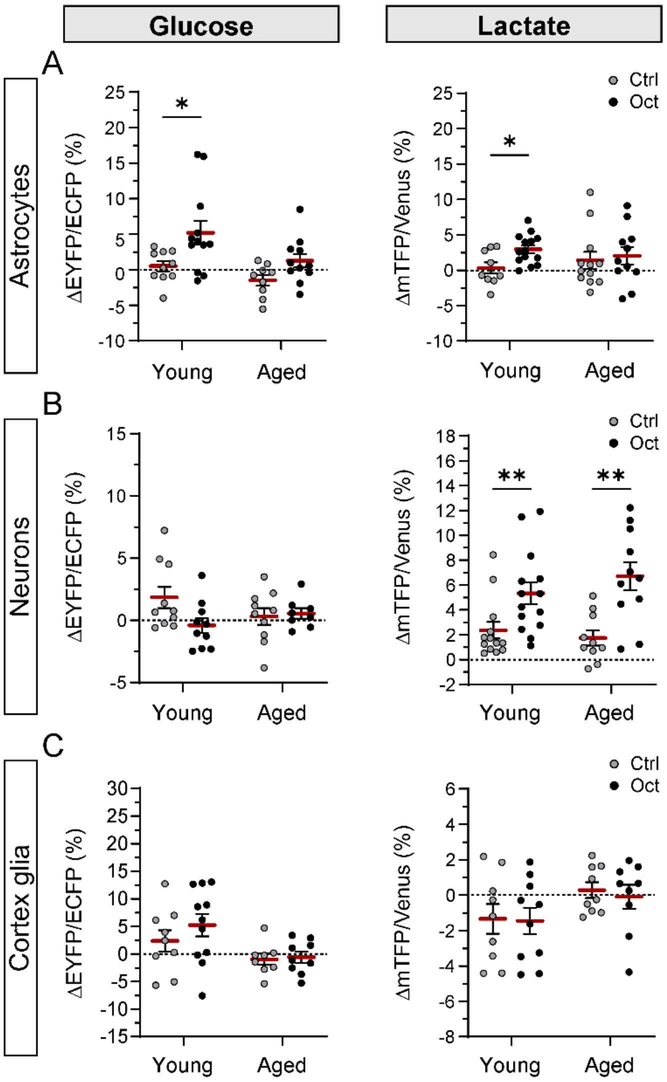


**Supplemental Fig. S6: Comparison of changes in cytosolic D-glucose and L-lactate levels in glial cells and neurons in brains of young and aged *Drosophila* upon vehicle versus octopamine stimulation.**

**A-C)** Scatter dot plots representing mean changes (in %) in the FLII$^{12}$Pglu-700µδ6 (ΔEYFP/ECFP; left panels) and Laconic (ΔmTFP/Venus; right panels) ratio signal after the addition of Ctrl (grey) and 10 µM Oct (black) in A) astrocytes, B) neurons and C) cortex glia in brains of young and aged *Drosophila*. Red lines represent mean of all data points; SEM is shown in black. Circles represent individual data points (mean values of all analyzed cells within individual brain/optic lobe). *$P < 0.05$, **$P < 0.01$ (Mann-Whitney *U*-test).

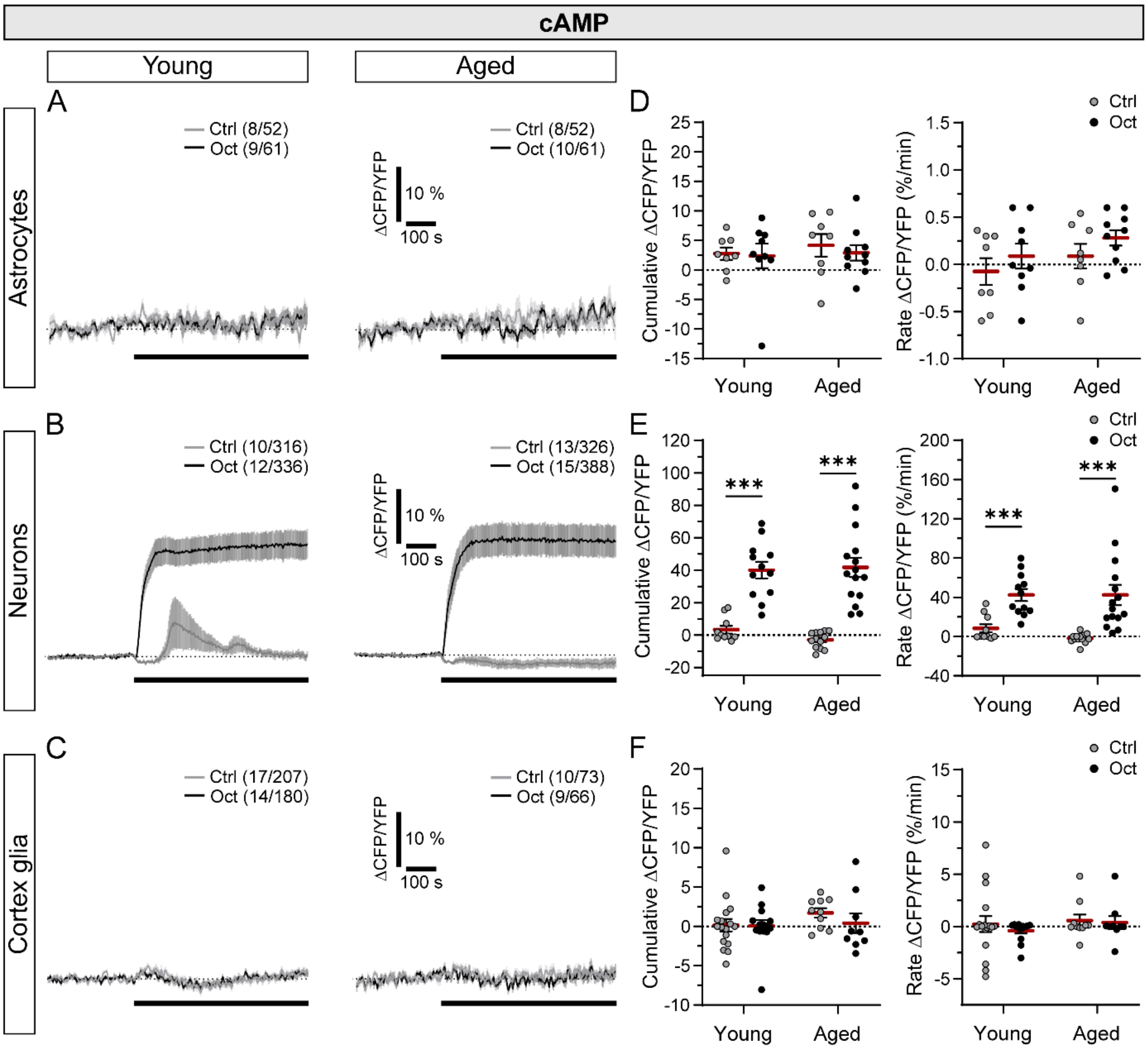


**Supplemental Fig. S7: Comparison of cytosolic cAMP signals in glial cells and neurons in brains of young and aged *Drosophila* upon vehicle versus octopamine stimulation.**

**A-C)** Mean time-dependent changes in the Epac1-camps fluorescence ratio signal (ΔCFP/YFP) upon the addition of experimental solution (vehicle; Ctrl; grey) and 10 µM octopamine (Oct; black) in A) astrocytes (*alrm>Epac1-camps*), B) neurons (*elav>Epac1-camps*), and C) cortex glia (*R55B12>Epac1-camps*) in brains of young (left panels) and aged (right panels) *Drosophila*. Data are expressed in percentages as a change in fluorescence ratio signal (ΔCFP/YFP) relative to the baseline values, reflecting changes in cytosolic concentration of cAMP. Numbers in parentheses depict the number of brains and cells analyzed. **D-F)** Scatter dot plots representing cumulative changes (Cumulative ΔCFP/YFP; left) and rate of changes in the Epac1-camps ratio signal (Rate ΔCFP/YFP (%/min); right) after the addition of Ctrl (grey) and 10 µM Oct (black) in D) astrocytes, E) neurons and F) cortex glia in brains of young and aged *Drosophila*. Red lines represent mean of all data points; SEM is shown in black. Circles represent individual data points (mean values of all analyzed cells within individual brain/optic lobe). ***$P < 0.001$ (Mann-Whitney *U*-test).

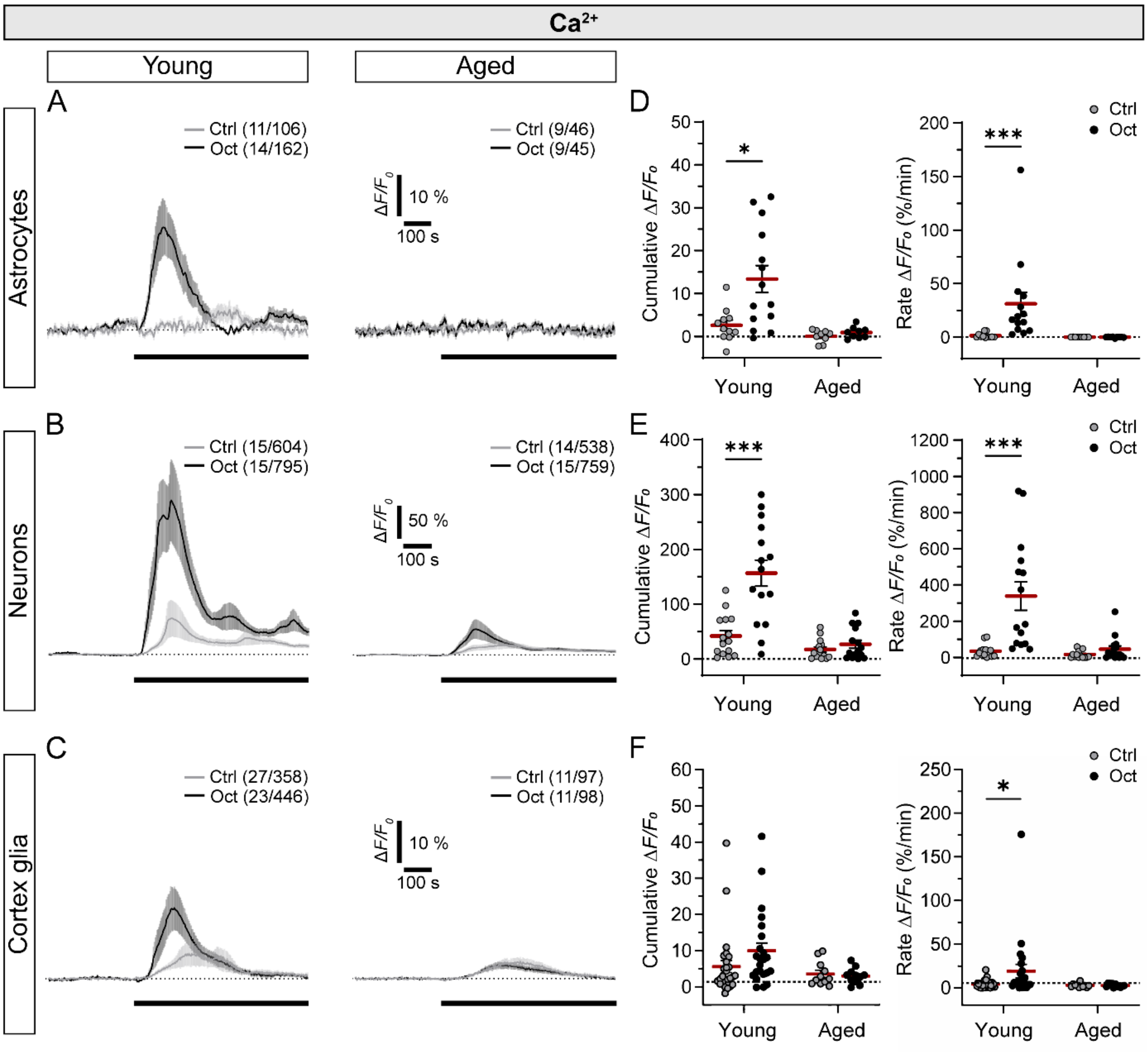


**Supplemental Fig. S8: Comparison of cytosolic $Ca^{2+}$ signals in glial cells and neurons in brains of young and aged *Drosophila* upon vehicle versus octopamine stimulation.**

**A-C)** Mean time-dependent changes in the jGCaMP7b fluorescence intensity signal ($\Delta F/F_0$) before and after the addition of experimental solution (vehicle; Ctrl; grey) and 10 µM octopamine (Oct; black) in A) astrocytes (*alrm>jGCaMP7b;mCherry*), B) neurons (*elav>jGCaMP7b;mCherry*), and C) cortex glia (*R55B12>jGCaMP7b;mCherry*) in brains of young (left panels) and aged (right panels) *Drosophila*. Data are expressed in percentages as a relative change in the fluorescence ($\Delta F/F_0$), reflecting changes in cytosolic concentration of $Ca^{2+}$. Numbers in parentheses depict the number of brains and cells analyzed. **D-F)** Scatter dot plots representing cumulative changes (Cumulative $\Delta F/F_0$; left) and rate of changes in the jGCaMP7b $\Delta F/F_0$ signal (Rate $\Delta F/F_0$ (%/min); right) after the addition of Ctrl (grey) and 10 µM Oct (black) in D) astrocytes, E) neurons and F) cortex glia in brains of young and aged *Drosophila*. Red lines represent mean of all data points; SEM is shown in black. Circles represent individual data points (mean values of all analyzed cells within individual brain/optic lobe). *$P$ < 0.05, ***$P$ < 0.001 (Unpaired t-test (D, left; F, right) or Mann-Whitney $U$-test (D, right; E)).

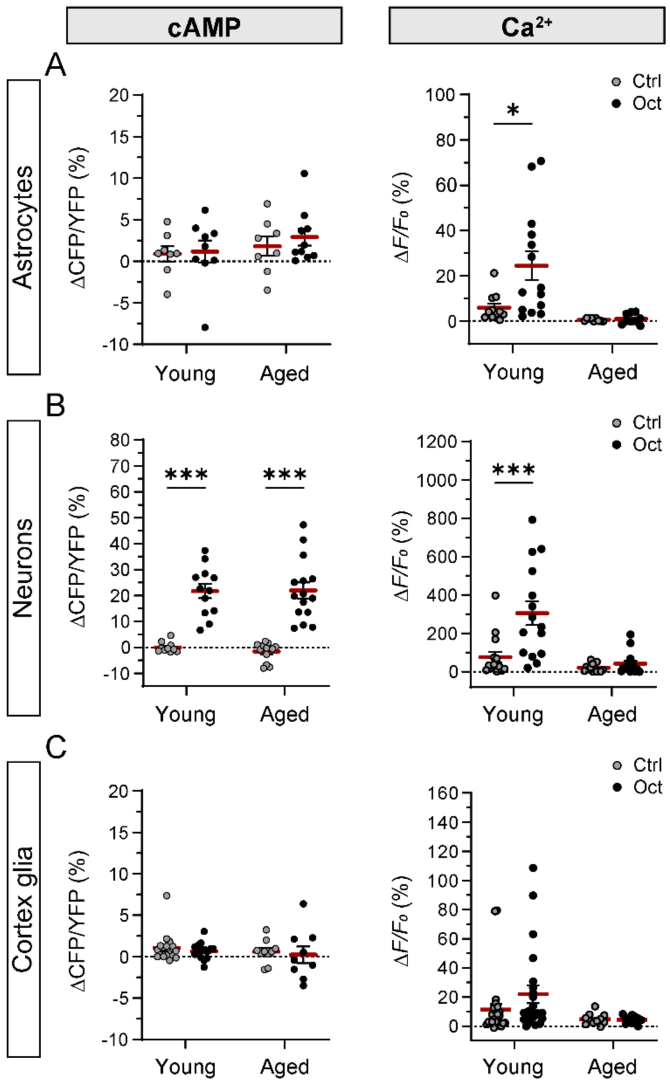


**Supplemental Fig. S9: Comparison of changes in cytosolic cAMP and $Ca^{2+}$ signals in glial cells and neurons in brains of young and aged *Drosophila* upon vehicle versus octopamine stimulation.**

**A-C)** Scatter dot plots representing mean changes (in %) in the Epac1-camps (ΔCFP/YFP; left panels) ratio signal and in the jGCaMP7b fluorescence intensity signal ($\Delta F/F_0$) after the addition of Ctrl (grey) and 10 µM Oct (black) in A) astrocytes, B) neurons and C) cortex glia in brains of young and aged *Drosophila*. Red lines represent mean of all data points; SEM is shown in black. Circles represent individual data points (mean values of all analyzed cells within individual brain/optic lobe). $^{*}P < 0.05$, $^{***}P < 0.001$ (Mann-Whitney *U*-test).

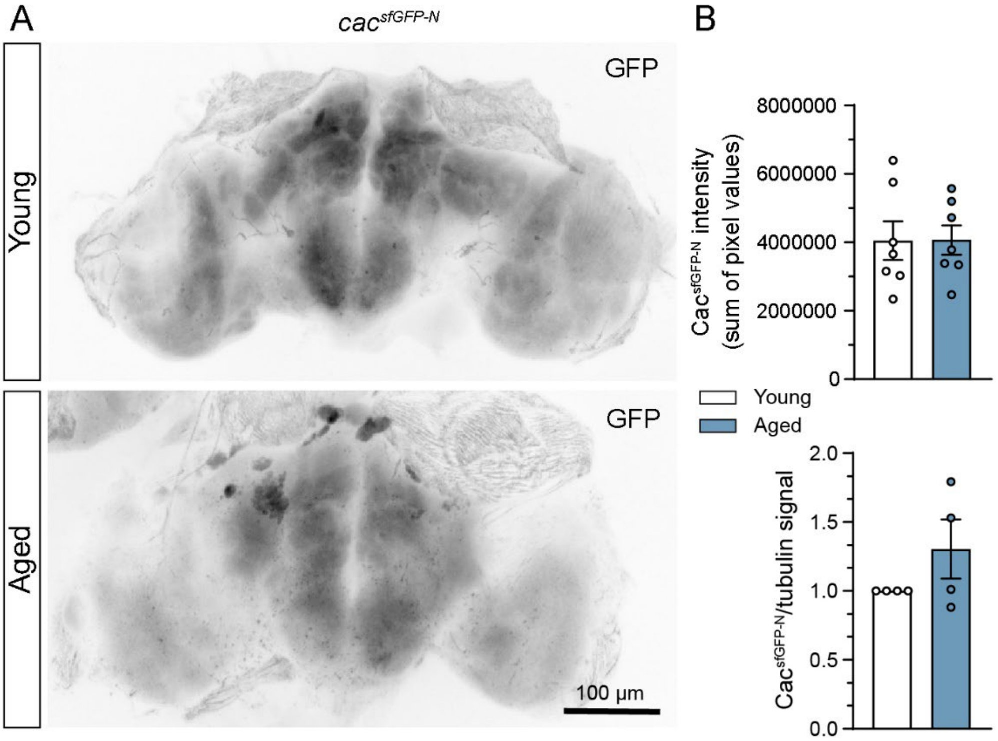


**Supplemental Fig. S10: Expression levels of the synaptic voltage-gated $Ca^{2+}$ channel Cacophony appears unaltered in aged brains.**

**A**) Representative images of fixed whole-brains from young (upper panels) and aged (lower panels) flies expressing superfold GFP-tagged voltage-gated $Ca^{2+}$ channel Cacophony under the native promotor (*Cacophony>GFP*). **B**) GFP intensity, quantified as the raw integrated density, calculated as the sum of the gray values of all pixels within the defined region (upper panel; n=7 brains) and WB analysis of the ratio between the sfGFP-tagged Cacophony and tubulin (lower pannel; n=4 samples, each sample consistent of 30 brains) in young (white) and aged (blue) *Drosophila*. Data are presented as means ± SEM. Circles represent individual data points.

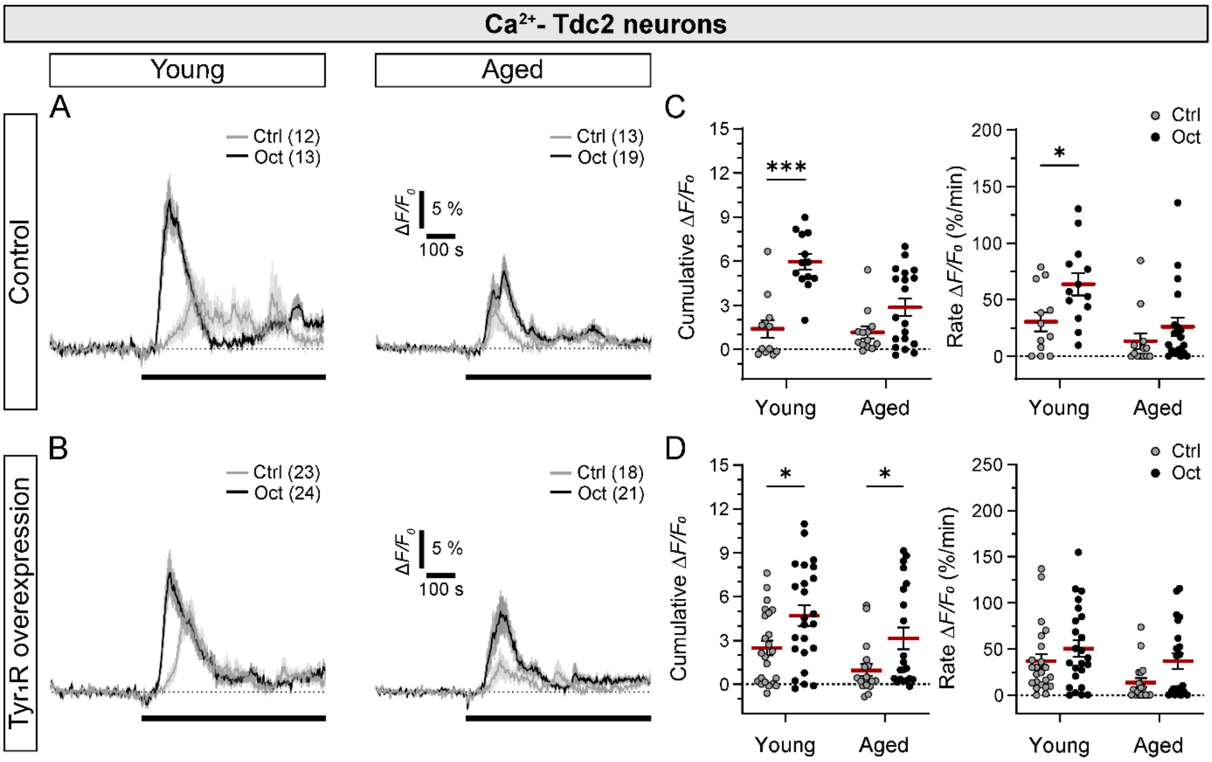


**Supplemental Fig. S11: Comparison of cytosolic $Ca^{2+}$ signals in Tdc2 neurons upon vehicle and octopamine stimulation in brains of young and aged control *Drosophila* and *Drosophila* overexpressing tyramine 1 receptors in Tdc2 neurons.**

**A-B)** Mean time-dependent changes in the jGCaMP7f fluorescence intensity signal ($\Delta F/F_0$) before and after the addition of experimental solution (vehicle; Ctrl; grey) and 10 µM octopamine (Oct; black) in octopaminergic/tyraminergic Tdc2+ neurons (Tdc2 neurons) in brains of young (left panels) and aged (right panels) *Drosophila* A) without (Control; *tdc2>jGCaMP7f;mCherry*) or B) with overexpression of tyramine 1 receptors in Tdc2 neurons ($Tyr_1R$ overexpression; *tdc2>jGCaMP7f;mCherry;Tyr*$_1$*R*). Data are expressed in percentages as a relative change in the fluorescence ($\Delta F/F_0$), reflecting changes in cytosolic concentration of $Ca^{2+}$. Numbers in parentheses depict the number of optic lobes analyzed. **C-D)** Scatter dot plots representing cumulative changes (Cumulative $\Delta F/F_0$; left) and rate of changes in the jGCaMP7f $\Delta F/F_0$ signal (Rate $\Delta F/F_0$ (%/min); right) after the addition of Ctrl (grey) and 10 µM Oct (black) in Tdc2 neurons in brains of young and aged *Drosophila* C) without (Control) or D) with overexpression of tyramine 1 receptors ($Tyr_1R$) in Tdc2 neurons ($Tyr_1R$ overexpression). Cumulative data is calculated within 3 min after stimulation. Red lines represent mean of all data points; SEM is shown in black. Circles represent individual data points (value per optic lobe). *$P < 0.05$, ***$P < 0.001$ (Mann-Whitney *U*-test).

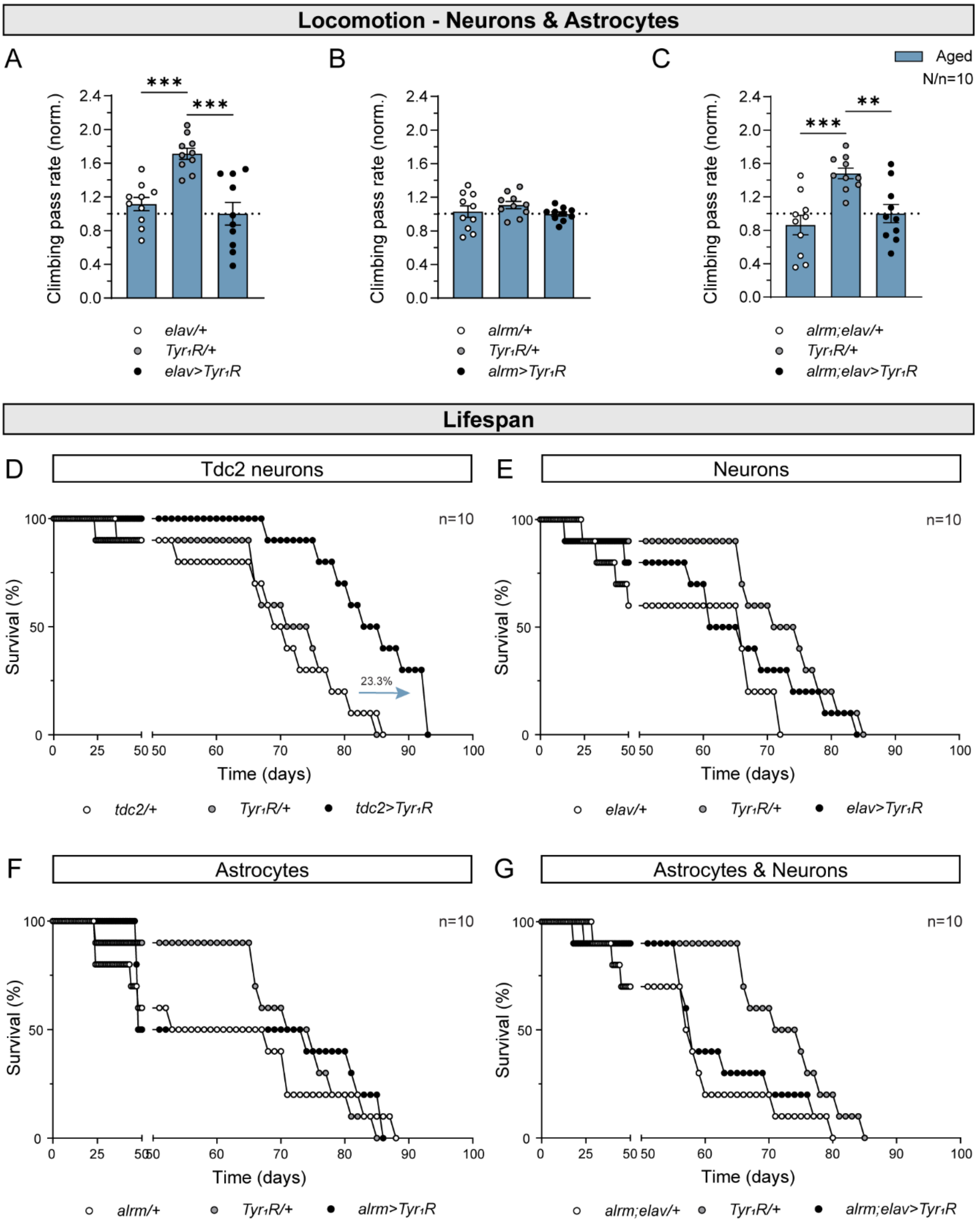


**Supplemental Fig. S12: Overexpression of tyramine 1 receptors in Tdc2 neurons extends lifespan of flies.**

**A-C)** Mean climbing pass rate of aged *Drosophila* without (*elav/+*, *alrm/+*, *alrm;elav/+* and *Tyr$_1$R/+*) or with overexpression of the tyramine 1 receptor (Tyr$_1$R) in neurons (*elav>Tyr$_1$R*), astrocytes (*alrm>Tyr$_1$R*) or both cell types (*alrm;elav>Tyr$_1$R*). N=number of independent vials (experiments), n=number of consecutive recordings per vial. Data are normalized to mean climbing pass rate of flies overexpressing $Tyr_1Rs$ within each data set and presented as means ± SEM. Circles represent individual data points (mean values per vial (independent experiment)). $^{**}P < 0.01$,$^{***}P < 0.001$ (one way ANOVA followed by Holm-Šídák's test). **D-G)** Lifespan assays of flies without (*tdc2/+*, *elav/+*, *alrm/+*, *alrm;elav/+* and *$Tyr_1R$/+*) or with overexpression of the $Tyr_1R$ in neurons (*elav>$Tyr_1R$*), astrocytes (*alrm>$Tyr_1R$*) or both cell types (*alrm;elav>$Tyr_1R$*; n=10 flies per genotype tested).

### Supplemental Tables

#### **Supplemental Table S1.** *Drosophila melanogaster* strains.

| Fly strain | Genotype | Additional information |
|---|---|---|
| *w1118* | *w1118;;* | - |
| *alrm-Gal4* | *w[*];alrm-Gal4;alrm-Gal4* | Gift from Christian Klämbt; Doherty, et al. [1] |
| *elav-Gal4* | *P{w[+mC]=GAL4-elav.L}2/CyO;* | BDSC #8765; Ogienko, et al. [2] |
| *R55B12-Gal4* | *w1118;;P{y[+t7.7]w[+mC]=GMR55B12-GAL4}attP2* | BDSC #39103; Jenett, et al. [3] |
| *R83E12-Gal4* | *w1118;;P{y[+t7.7] w[+mC]=GMR83E12-GAL4}attP2* | BDSC #40363; Jenett, et al. [3] |
| *R85G01-Gal4* | *w1118;;P{y[+t7.7] w[+mC]=GMR85G01-GAL4}attP2* | BDSC #40436; Jenett, et al. [3] |
| *SPG-Gal4* | *w[*]; P{w[+m*]=moody-GAL4.SPG}2;* | BDSC #90883; Padiath, et al. [4] |
| *Oct-TyrR-2A-Gal4* | *TI{RFP[3xP3.cUa]=2A-GAL4}Oct-TyrR[2A-GAL4.KI]* | BDSC #86138 |
| *tdc2-Gal4* | *;;tdc2-Gal4 w+* | Gift from Mareike Selcho; Cole, et al. [5] |
| *UAS-jGCaMP7b* | *w1118;P{20xUAS-IVS-jGCaMP7b}su(Hw)attP5;* | BDSC #80907; Dana, et al. [6] |
| *UAS-mCherry* | *;;20XUAS-6XmCherry-HA}attP2w+/TM6B,Tb* | BDSC #52268 |
| *UAS-Epac1-camps* | *w1118;;UAS-Epac1-camps* $w^{+}$*/Sb* | Maiellaro, et al. [7] |
| *UAS-FLII*$^{12}$*Pglu-700μδ6* | *w1118;;pUASt-FLII*$^{12}$*Pglu-700μδ6 w+/Cyo* | Volkenhoff, et al. [8] |
| *UAS-Laconic* | *w1118;;pUASt-Laconic w+/Cyo* | This study. |
| *UAS-TransTimer[sfGFP,RFP]* | *v[*]; If/CyO; UAS-TransTimer [v+]attP2* | Gift from Richard Binari; He, et al. [9] |
| *UAS-Oct-TyrR* | *w[*];P{y[=t7.7]w[+mC]=UAS-Oct-TyrR}attP40* | BDSC #95289 |
| *UAS-jGCaMP7f* | *w1118;;{20xUAS-IVS-jGCaMP7f}VK00005w+* | BDSC #97031; Dana, et al. [6] |
| *Cac*$^{sfGFP\text{-}N}$ | | Gratz, et al. [10] |

BDSC, Bloomington *Drosophila* Stock Center (USA); Chr, chromosome.

**Supplemental Table S2.** Mean vacuole number, individual vacuole area and total vacuole area per brain and mean climbing pass rate of young and aged *Drosophila*.

| **Neurodegeneration parameter** | **Young** | **Aged** | **n** |
|---|---|---|---|
| No. of vacuoles | 3.09 ± 0.73 | 5.27 ± 0.65# | 11 |
| Individual vacuole area (μm$^2$) | 7.64 ± 1.28 | 9.45 ± 0.65# | 11 |
| Total vacuole area (μm$^2$) | 22.45 ± 5.17 | 52.99 ± 9.12## | 11 |
| **Locomotion performance** | | | **N/n** |
| Climbing pass rate (%) | 88.12 ± 2.56 | 12.30 ± 2.81### | 10/10 |

n, number of brains analysed; N/n, number of independent vials (experiments)/number of consecutive recordings per vial. Data are presented as means ± standard error of the mean (related to Figure 1).

##*P* < 0.01; ###*P* < 0.001 versus Young (Student's t-test); #*P* < 0.05 versus Young (Mann-Whitney *U*-test).

**Supplemental Table S3.** Mean basal FLII[12]Pglu-700µδ6 and Laconic fluorescence ratio signal values normalized to maximal response upon stimulation with 20 mM D-glucose and 30 mM L-lactate, respectively, in astrocytes, cortex glia and neurons in brains isolated from young and aged *Drosophila*.

| | Astrocytes | *n* | Cortex glia | *n* | Neurons | *n* |
|---|---|---|---|---|---|---|
| **FLII[12]Pglu-700µδ6 ratio (EYFP/ECFP)** | | | | | | |
| **Young** | 0.90 ± 0.02 | 10 | 0.86 ± 0.02 | 11 | 0.86 ± 0.01 | 12 |
| **Aged** | 0.95 ± 0.01 | 9 | 0.89 ± 0.02 | 11 | 0.89 ± 0.00$ | 9 |
| **Laconic ratio (mTFP/Venus)** | | | | | | |
| **Young** | 0.84 ± 0.01 | 13 | 0.85 ± 0.01 | 10 | 0.87 ± 0.01 | 14 |
| **Aged** | 0.85 ± 0.01 | 12 | 0.81 ± 0.01# | 11 | 0.87 ± 0.01*** | 15 |

EYFP/ECFP, FLII[12]Pglu-700µδ6 fluorescence ratio signal; FLII[12]Pglu-700µδ6, genetically encoded glucose sensor; mTFP/Venus, Laconic fluorescence ratio signal; Laconic, genetically encoded lactate sensor; n, number of brains analysed. Data are presented as means ± standard error of the mean (related to Figure 2 and Supplemental Figure S2).

#$P < 0.05$ versus Young (Unpaired t-test).
$ $P < 0.05$ versus astrocytes (ANOVA on ranks, followed by Dunn's test).
***$P < 0.001$ versus cortex glia (Brown-Forsythe and Welch ANOVA, followed by Dunnett's T3 test).

**Supplemental Table S4.** Mean cumulative changes and rate of changes in the **FLII$^{12}$Pglu-700µδ6** fluorescence ratio signal in astrocytes, neurons and cortex glia in brains isolated from young and aged *Drosophila* on stimulation with control solution, octopamine and high extracellular D-glucose concentration.

| | **Cumulative ΔEYFP/ECFP** | | **Rate ΔEYFP/ECFP (%/min)** | | ***n/N*** | |
|---|---|---|---|---|---|---|
| | **Young** | **Aged** | **Young** | **Aged** | **Young** | **Aged** |
| **Astrocytes** | | | | | | |
| Control | 0.61 ± 0.31 | 0.03 ± 0.19 | 0.13 ± 0.21 | -0.14 ± 0.18 | 10/71 | 9/58 |
| Oct | 2.82 ± 0.82* | 0.50 ± 0.40# | 4.21 ± 1.33* | -0.08 ± 0.20# | 12/68 | 11/73 |
| Glc | 7.64 ± 0.72 | 8.20 ± 1.11 | 9.72 ± 1.64 | 5.53 ± 1.04 | 10/66 | 9/48 |
| **Neurons** | | | | | | |
| Control | 0.34 ± 0.30 | 0.15 ± 0.20 | -0.02 ± 0.10 | 0.02 ± 0.07 | 10/132 | 10/199 |
| Oct | -0.19 ± 0.30 | 0.17 ± 0.18 | -0.50 ± 0.68 | 0.01 ± 0.07 | 10/155 | 8/226 |
| Glc | 10.92 ± 1.08 | 6.36 ± 0.32## | 15.90 ± 1.39 | 8.73 ± 1.21## | 12/228 | 9/219 |
| **Cortex glia** | | | | | | |
| Control | 1.83 ± 0.98 | -0.13 ± 0.66 | 2.69 ± 1.81 | -0.09 ± 0.09 | 9/78 | 8/58 |
| Oct | 3.66 ± 0.94 | -0.18 ± 0.56 | 4.43 ± 2.12 | -0.12 ± 0.10 | 11/112 | 9/58 |
| Glc | 10.41 ± 0.94 | 8.55 ± 1.80 | 9.00 ± 0.87 | 9.11 ± 1.49 | 11/110 | 11/67 |

ΔEYFP/ECFP, change in the normalized FLII$^{12}$Pglu-700µδ6 fluorescence ratio signal; FLII$^{12}$Pglu-700µδ6, genetically encoded glucose sensor; *n*, number of brains analysed; *N*, number of cells; Control, control solution (vehicle); Oct, octopamine (10 µM); Glc, D-glucose (20 mM). Data are presented as means ± standard error of the mean (related to Figures 2 and 3, Supplemental Figures S1 and S3).

##*P* < 0.01 versus Young (Student's t-test); #*P* < 0.05 versus Young (Mann-Whitney *U*-test).
**P* < 0.05 versus Control (Mann-Whitney *U*-test).

**Supplemental Table S5.** Mean cumulative changes and rate of changes in the **Laconic** fluorescence ratio signal in astrocytes, neurons and cortex glia in brains isolated from young and aged *Drosophila* on stimulation with control solution, octopamine, high extracellular L-lactate concentration and α-cyano-4-hydroxycinnamate.

| | **Cumulative ΔmTFP/Venus** | | **Rate ΔmTFP/Venus (%/min)** | | ***n*/*N*** | |
|---|---|---|---|---|---|---|
| | **Young** | **Aged** | **Young** | **Aged** | **Young** | **Aged** |
| **Astrocytes** | | | | | | |
| Control | 0.61 ± 0.37 | 0.46 ± 0.39 | 0.06 ± 0.03 | 0.09 ± 0.09 | 9/74 | 12/55 |
| Oct | 0.74 ± 0.20*** | 0.29 ± 0.30 | 0.34 ± 0.05 | 0.28 ± 0.10 | 14/133 | 11/63 |
| Lac | 12.19 ± 1.00 | 10.68 ± 0.90 | 28.48 ± 4.51 | 23.10 ± 4.14 | 13/91 | 12/53 |
| CHC | - | - | 2.10 ± 0.55 | - | 6/74 | - |
| **Neurons** | | | | | | |
| Control | 1.31 ± 0.34 | 1.05 ± 0.34 | 3.31 ± 1.02 | 2.13 ± 1.03 | 13/424 | 10/270 |
| Oct | 2.49 ± 0.40* | 2.95 ± 0.58* | 3.71 ± 0.66 | 4.33 ± 1.17 | 14/467 | 11/326 |
| Lac | 8.91 ± 0.71 | 8.69 ± 0.60 | 21.43 ± 2.56 | 21.52 ± 2.77 | 14/325 | 15/386 |
| CHC | - | - | 0.67 ± 0.25[&] | - | 7/111 | - |
| **Cortex glia** | | | | | | |
| Control | -0.95 ± 0.30 | -0.15 ± 0.21[#] | -0.51 ± 0.30 | 0.02 ± 0.05 | 9/70 | 9/50 |
| Oct | -1.09 ± 0.23[$$$] | -0.22 ± 0.25 | -0.60 ± 0.36[$$$] | 0.00 ± 0.07 | 10/75 | 9/62 |
| Lac | 10.16 ± 0.99 | 11.73 ± 0.93 | 23.46 ± 1.96 | 25.47 ± 7.37 | 10/105 | 11/70 |
| **Neuropil EG** | | | | | | |
| Oct | -0.21 ± 0.16[$$$] | - | -0.12 ± 0.06[$$$] | - | 11/76 | - |
| **Perineurial glia** | | | | | | |
| Oct | 0.44 ± 0.12[$] | - | 0.39 ± 0.17[$$] | - | 13[a] | - |
| **Subperineurial glia** | | | | | | |
| Oct | 0.64 ± 0.42[$$] | - | 0.33 ± 0.21[$$$] | - | 12[a] | - |

ΔmTFP/Venus, change in the normalized Laconic fluorescence ratio signal; Laconic, genetically encoded lactate sensor; *n*, number of brains analysed; *N*, number of cells; EG, ensheathing glia; Control, control solution (vehicle); Oct, octopamine (10 µM); Lac, L-lactate (30 mM); CHC, α-cyano-4-hydroxycinnamate (6 mM). Data are presented as means ± standard error of the mean (related to Figures 2 and 4, Supplemental Figures S1, S4 and S5).

[a]Due to the native distribution of cell types within the *Drosophila* brain, the analysis was carried out using one or more ROIs covering larger brain areas per individual brain.

[#]*P* < 0.05 versus Young (Student's t-test).
****P* < 0.001 versus Control (Student's t-test); **P* < 0.05 versus Control (Mann-Whitney *U*-test).
[&]*P* < 0.05 versus astrocytes (Mann-Whitney *U*-test).

$^{\$}P < 0.05$, $^{\$\$}P < 0.01$, $^{\$\$\$}P < 0.001$ versus Neurons (One-way ANOVA on Ranks, followed by Dunn’s test)

**Supplemental Table S6**. Mean cumulative changes and rate of changes in the **Epac1-camps** fluorescence ratio signal in astrocytes, neurons and cortex glia in brains isolated from young and aged Drosophila on stimulation with control solution and octopamine.

| | Cumulative ΔCFP/YFP | | Rate ΔCFP/YFP (%/min) | | *n*/*N* | |
|---|---|---|---|---|---|---|
| | **Young** | **Aged** | **Young** | **Aged** | **Young** | **Aged** |
| **Astrocytes** | | | | | | |
| Control | 2.76 ± 1.03 | 4.18 ± 1.91 | -0.08 ± 0.14 | 0.09 ± 0.13 | 8/52 | 8/52 |
| Oct | 2.40 ± 2.07 | 2.91 ± 1.31 | 0.09 ± 0.13 | 0.28 ± 0.08 | 9/61 | 10/61 |
| **Neurons** | | | | | | |
| Control | 3.29 ± 2.34 | -3.01 ± 1.40 | 8.45 ± 4.11 | -1.63 ± 1.34# | 10/316 | 13/326 |
| Oct | 40.05 ± 5.05*** | 41.72 ± 6.03*** | 42.40 ± 6.13*** | 42.40 ± 10.27*** | 12/336 | 15/388 |
| **Cortex glia** | | | | | | |
| Control | 0.15 ± 0.79 | 1.70 ± 0.60 | 0.24 ± 0.77 | 0.56 ± 0.57 | 17/207 | 10/73 |
| Oct | 0.07 ± 0.75 | 0.41 ± 1.25 | -0.41 ± 0.25 | 0.37 ± 0.64 | 14/180 | 9/66 |

ΔCFP/YFP, change in the normalized Epac1-camps fluorescence ratio signal; Epac1-camps, genetically encoded cAMP sensor; *n*, number of brains analysed; *N*, number of cells; Control, control solution (vehicle); Oct, octopamine (10 µM). Data are presented as means ± standard error of the mean (related to Figure 5, Supplemental Figure S7).

#$P < 0.05$ versus Young (Mann-Whitney *U*-test).
***$P < 0.001$ versus Control (Mann-Whitney *U*-test).

**Supplemental Table S7.** Mean cumulative changes and rate of changes in the **jGCaMP7b** fluorescence signal in astrocytes, neurons and cortex glia in brains isolated from young and aged *Drosophila* on stimulation with control solution, octopamine, thapsigargin and ionomycin.

| | Cumulative $\Delta F/F_0$ | | Rate $\Delta F/F_0$ (%/min) | | $n/N$ | |
|---|---|---|---|---|---|---|
| | **Young** | **Aged** | **Young** | **Aged** | **Young** | **Aged** |
| **Astrocytes** | | | | | | |
| Control | 2.63 ± 1.18 | 0.07 ± 0.48 | 1.36 ± 0.73 | -0.07 ± 0.04 | 11/106 | 9/46 |
| Oct | 13.38 ± 3.14* | 0.90 ± 0.44## | 31.16 ± 10.74*** | -0.16 ± 0.14### | 14/162 | 9/45 |
| **Neurons** | | | | | | |
| Control | 41.94 ± 9.74 | 16.97 ± 4.88# | 34.10 ± 8.84 | 16.78 ± 5.60 | 15/604 | 14/538 |
| Oct | 156.50 ± 23.67*** | 26.65 ± 7.21### | 338.90 ± 78.08*** | 45.84 ± 17.33### | 15/795 | 15/759 |
| TG | 88.43 ± 11.40 | 10.79 ± 2.39### | 121.40 ± 21.09 | 7.63 ± 1.15## | 8/294 | 7/220 |
| Iono | 244.60 ± 32.25 | 85.33 ± 15.38### | 201.90 ± 54.39 | 29.62 ± 7.53### | 11/668 | 13/637 |
| **Cortex glia** | | | | | | |
| Control | 5.58 ± 1.68 | 3.61 ± 1.02 | 4.22 ± 0.87 | 2.69 ± 0.66 | 27/358 | 11/97 |
| Oct | 9.97 ± 2.17* | 2.98 ± 0.66 | 19.10 ± 7.65* | 2.32 ± 0.55 | 23/446 | 11/98 |

*F*, fluorescence; $\Delta F/F_0$, change in the normalized jGCaMP7b fluorescence signal; jGCaMP7b, genetically encoded $Ca^{2+}$ sensor; *n*, number of brains analysed; *N*, number of cells; Control, control solution (vehicle); Oct, octopamine (10 µM); TG, thapsigargin (1 µM); Iono, ionomycin (10 µM). Data are presented as means ± standard error of the mean (related to Figure 6, Supplemental Figure S8).

$^{\#\#}P < 0.01$, $^{\#\#\#}P < 0.001$ versus Young (Student's t-test); $^{\#}P < 0.05$, $^{\#\#\#}P < 0.001$ versus Young (Mann-Whitney *U*-test).
$^{*}P < 0.05$ versus Control (Student's t-test); $^{*}P < 0.05$, $^{***}P < 0.001$ versus Control (Mann-Whitney *U*-test).

**Supplemental Table S8**. Mean changes in the **FLII$^{12}$Pglu-700µδ6**, **Laconic** and **Epac1-camps** fluorescence ratio signal and in the **jGCaMP7b** fluorescence signal in astrocytes, neurons and cortex glia in brains isolated from young and aged *Drosophila* on stimulation with control solution and octopamine.

| | **FLII$^{12}$Pglu-700µδ6 ΔEYFP/ECFP (%)** | | **Laconic ΔmTFP/Venus (%)** | | **Epac1-camps ΔCFP/YFP (%)** | | **jGCaMP7b Δ*F*/*F*$_0$ (%)** | |
|---|---|---|---|---|---|---|---|---|
| | **Young** | **Aged** | **Young** | **Aged** | **Young** | **Aged** | **Young** | **Aged** |
| **Astrocytes** | | | | | | | | |
| Control | 0.50 ± 0.71 | -1.51 ± 0.77 | 0.31 ± 0.81 | 1.40 ± 1.20 | 0.87 ± 0.92 | 1.81 ± 1.17 | 5.93 ± 1.82 | 0.56 ± 0.22 |
| Oct | 5.19 ± 1.69* | 1.25 ± 0.97 | 2.95 ± 0.56* | 2.01 ± 1.25 | 1.17 ± 1.34 | 2.92 ± 1.02 | 24.43 ± 6.27* | 0.95 ± 0.79 |
| **Neurons** | | | | | | | | |
| Control | 1.85 ± 0.86 | 0.32 ± 0.67 | 2.36 ± 0.67 | 1.74 ± 0.61 | 0.05 ± 0.64 | -1.59 ± 0.99 | 76.48 ± 27.59 | 21.45 ± 5.53 |
| Oct | -0.40 ± 0.61 | 0.57 ± 0.42 | 5.35 ± 0.89** | 6.71 ± 1.14** | 21.70 ± 2.78*** | 21.95 ± 3.10*** | 305.80 ± 62.62*** | 42.48 ± 14.60 |
| **Cortex glia** | | | | | | | | |
| Control | 2.37 ± 1.96 | -0.95 ± 1.04 | -1.34 ± 0.84 | 0.28 ± 0.44 | 1.07 ± 0.43 | 0.60 ± 0.45 | 11.28 ± 3.88 | 4.69 ± 1.16 |
| Oct | 5.25 ± 2.06 | -0.60 ± 1.00 | -1.46 ± 0.74 | -0.09 ± 0.68 | 0.68 ± 0.28 | 0.24 ± 1.01 | 21.92 ± 6.02 | 4.28 ± 0.82 |

ΔEYFP/ECFP, change in the normalized FLII$^{12}$Pglu-700µδ6 fluorescence ratio signal; FLII$^{12}$Pglu-700µδ6, genetically encoded glucose sensor; ΔmTFP/Venus, change in the normalized Laconic fluorescence ratio signal; Laconic, genetically encoded lactate sensor; ΔCFP/YFP, change in the normalized Epac1-camps fluorescence ratio signal; Epac1-camps, genetically encoded cAMP sensor; *F*, fluorescence; $\Delta F/F_0$, change in the normalized jGCaMP7b fluorescence signal; jGCaMP7b, genetically encoded $Ca^{2+}$ sensor; Control, control solution (vehicle); Oct, octopamine (10 µM). Data are presented as means ± standard error of the mean (related to Supplemental Figures S6 and S9).

#$P < 0.05$, **$P < 0.01$, ***$P < 0.001$ versus Control (Mann-Whitney *U*-test).

**Supplemental Table S9.** Mean sum of GFP-positive pixels reflecting synaptic voltage-gated $Ca^{2+}$ channel Cacophony expression and ratio of GFP-tagged Cacophony and tubulin in young and aged *Drosophila*.

| | **Young** | **Aged** | **n** |
|---|---|---|---|
| Cac::GFP Intensity (sum of all pixels) | 4,050,569.00 ± 563,730.00 | 4,068,246.00 ± 425,417.00 | 7 |
| Cac::GFP/tubulin ratio | 1.00 ± 0.00 | 1.31 ± 0.21 | 4 |

Cac, Cacophony; GFP, green fluorescent protein; n, number of brains/samples analysed. Data are presented as means ± standard error of the mean (related to Supplemental Figure S10).

**Supplemental Table S10.** Mean changes, cumulative changes and rate of changes in the **jGCaMP7f** fluorescence signal in Tdc2 neurons in brains isolated from young and aged *Drosophila* in control and flies overexpressing tyramine 1 receptors on stimulation with control solution and octopamine.

| | **$\Delta F/F_0$ (%)[a]** | | **Cumulative $\Delta F/F_0$[a]** | | **Rate $\Delta F/F_0$ (%/min)** | | ***n*** | |
|---|---|---|---|---|---|---|---|---|
| | **Young** | **Aged** | **Young** | **Aged** | **Young** | **Aged** | **Young** | **Aged** |
| **Control** | | | | | | | | |
| Control | 10.80 ± 4.06 | 8.56 ± 3.04 | 1.39 ± 0.61 | 1.16 ± 0.41 | 30.53 ± 8.43 | 13.12 ± 6.90 | 12 | 13 |
| Oct | 29.70 ± 3.04*** | 15.44 ± 2.92## | 5.95 ± 0.53*** | 2.86 ± 0.58## | 63.60 ± 9.79* | 26.06 ± 8.12## | 13 | 19 |
| **$Tyr_1R$ overexpression** | | | | | | | | |
| Control | 16.63 ± 2.75 | 7.09 ± 2.17# | 2.49 ± 0.52 | 0.97 ± 0.42 | 36.70 ± 7.74 | 13.38 ± 4.89## | 23 | 18 |
| Oct | 23.24 ± 3.34 | 17.57 ± 3.85 | 4.71 ± 0.69* | 3.15 ± 0.75* | 50.56 ± 8.79 | 36.87 ± 8.73 | 24 | 21 |

*F*, fluorescence; $\Delta F/F_0$, change in the normalized jGCaMP7f fluorescence signal; jGCaMP7f, genetically encoded $Ca^{2+}$ sensor; *n*, number of regions(optic lobes) analysed; Control, control solution (vehicle); Oct, octopamine (10 µM); $Tyr_1R$, tyramine 1 receptor. Data are presented as means ± standard error of the mean (related to Figure 7, Supplemental Figure S11).

[a]$\Delta F/F_0$ and Cumulative $\Delta F/F_0$ were calculated within 3 min after the addition of stimulus.

#$P < 0.05$; ##$P < 0.01$ versus Young (Mann-Whitney *U*-test).
*$P < 0.05$, ***$P < 0.001$ versus Control (Mann-Whitney *U*-test).

**Supplemental Table S11.** Mean endogenous tyramine 1 receptor expression level in brains of young and aged *Drosophila* and mean climbing pass rate of aged *Drosophila* without or with overexpression of the tyramine 1 receptor in Tdc2 neurons.

| **$Tyr_1R$ expression** | **Young** | **Aged** | **n** |
|---|---|---|---|
| Mean sfGFP/RFP intensity | 0.89 ± 0.06 | 0.60 ± 0.04$^{\#\#}$ | 11/10 |
| **Climbing pass rate (%)** | - | **Aged** | **N/n** |
| *tdc2/+* | - | 48.10 ± 5.86$^{\$}$ | 10/10 |
| *$Tyr_1R$/+* | - | 29.20 ± 7.45$^{\$\$\$}$ | 10/10 |
| *tdc2>$Tyr_1R$* | - | 84.50 ± 4.39 | 10/10 |

$Tyr_1R$, tyramine 1 receptor; sfGFP, fast-folding green fluorescent protein; RFP, red fluorescent protein; n, number of brains analysed; tdc2, tyrosine decarboxylase 2; N/n, number of independent vials (experiments)/number of consecutive recordings per vial. Data are presented as means ± standard error of the mean (related to Figure 7).

$^{\#\#}P < 0.05$ versus Young (Mann-Whitney *U*-test).
$^{\$}P < 0.05$, $^{\$\$\$}P < 0.001$ versus *tdc2>$Tyr_1R$* (One-way ANOVA on Ranks, followed by Dunn's test)

**Supplemental Table S12.** Mean climbing pass rate of young and aged Drosophila without or with overexpression of the tyramine 1 receptor in neurons, astrocytes or both cell types.

| **Climbing pass rate (%)** | **Young** | **Aged** | **N/n** |
|---|---|---|---|
| *elav/+* | 92.6 ± 1.18 | 40.8 ± 2.91 | 10/10 |
| *$Tyr_1R$/+* | 97.4 ± 1.10 | 62.7 ± 2.42 | 10/10 |
| *elav>$Tyr_1R$* | 97.8 ± 0.80 | 36.6 ± 4.90 | 10/10 |
| *alrm/+* | 95.9 ± 0.66 | 58.2 ± 3.91 | 10/10 |
| *alrm>$Tyr_1R$* | 94.0 ± 0.96 | 56.6 ± 1.45 | 10/10 |
| *alrm;elav/+* | 96.9 ± 0.80 | 31.4 ± 4.26 | 10/10 |
| *alrm;elav>$Tyr_1R$* | 92.6 ± 1.52 | 36.4 ± 3.98 | 10/10 |

$Tyr_1R$, tyramine 1 receptor; N/n, number of independent vials (experiments)/number of consecutive recordings per vial. Data are presented as means ± standard error of the mean (related to Supplemental Figure S12).

**Supplemental Table S13.** Mean lifespan without or with overexpression of the tyramine 1 receptor in Tdc2 neurons, neurons, astrocytes or both cell types.

| | Mean lifespan (days) | n |
|---|---|---|
| *tdc2/+* | 66.90 ± 4.41 | 10 |
| *$Tyr_1R$/+* | 83.10 ± 2.64 | 10 |
| *elav/+* | 54.90 ± 5.54 | 10 |
| *$Tyr_1R$/+* | 67.90 ± 5.39 | 10 |
| *elav>$Tyr_1R$* | 60.40 ± 6.23 | 10 |
| *alrm/+* | 58.60 ± 6.41 | 10 |
| *alrm>$Tyr_1R$* | 63.70 ± 5.80 | 10 |
| *alrm;elav/+* | 54.70 ± 4.70 | 10 |
| *alrm;elav>$Tyr_1R$* | 59.10 ± 5.42 | 10 |

$Tyr_1R$, tyramine 1 receptor; tdc2, tyrosine decarboxylase 2; n, number of flies. Data are presented as means ± standard error of the mean (related to Supplemental Figure S12).
